# Theoretical Guarantees for Phylogeny Inference from Single-Cell Lineage Tracing

**DOI:** 10.1101/2021.11.21.469464

**Authors:** Robert Wang, Richard Zhang, Alex Khodaverdian, Nir Yosef

**Author notes:** Equal contribution (Robert Wang, Richard Zhang, Alex Khodaverdian).

## Abstract

CRISPR-Cas9 lineage tracing technologies have emerged as a powerful tool for investigating development in single-cell contexts, but exact reconstruction of the underlying clonal relationships in experiment is plagued by data-related complications. These complications are functions of the experimental parameters in these systems, such as the Cas9 cutting rate, the diversity of indel outcomes, and the rate of missing data. In this paper, we develop two theoretically grounded algorithms for reconstruction of the underlying phylogenetic tree, as well as asymptotic bounds for the number of recording sites necessary for exact recapitulation of the ground truth phylogeny at high probability. In doing so, we explore the relationship between the problem difficulty and the experimental parameters, with implications for experimental design. Lastly, we provide simulations validating these bounds and showing the empirical performance of these algorithms. Overall, this work provides a first theoretical analysis of phylogenetic reconstruction in the CRISPR-Cas9 lineage tracing technology.

## Introduction

Phylogenetic trees are routinely constructed to describe the developmental relationships within sets of extant taxa such as different organisms, proteins, or single cells. A landmark early example of using phylogenetics to describe cellular relationships was that of Sulston and colleagues reporting the development of C. elegans as deduced from meticulous visual observation [1, 2]. Recent progress in CRISPR-Cas9 based lineage tracing technologies now enables the inference of cellular lineage relationships in more complex organisms where visual observation is not possible. This is owing CRISPR-Cas9’s ability to generate heritable, irreversible, and information-rich mutation events that can be read through single-cell assays (such as RNA-seq) and subsequently used to infer the underlying phylogeny [3, 4, 5, 6, 7, 8, 9, 10, 11, 12]. Typically, these technologies start by engineering a single progenitor cell with artificial transcribed recording sites that accumulate stable insertions or deletions (“indels”) as a result of repair of Cas9 double-stranded breaks. These indel mutations are subsequently inherited by future descendants, and the accumulation of these mutations is used to infer the clonal relationships between the observed cells, stratifying them into clades of increasing resolution. Thus far, studies have paired these technologies with single-cell transcriptomic profiling [13] to study questions in development (e.g., inferring lineage relationships between cellular compartments) [7, 8, 6, 5, 9, 11] and cancer progression (e.g., inferring rates and routes of metastases) [12, 14].

Despite the many advantages of CRISPR-Cas9 lineage tracing systems, an outstanding goal is develop methods to accurately infer the underlying developmental process and to determine under what experimental conditions the problem is tractable. Exact reconstruction of the ground truth phylogenetic tree, defined here as having a reconstructed tree that exactly matches that of the ground truth clonal relationships, is plagued by various data-related complications (Fig.1). First, convergent evolution (or homoplasy) events can occur whereby the cells might appear to be incorrectly related to each other because the same indel occurs in unrelated clades. [15, 16, 10]. Second, substantial missing data is observed in these experiments in which the information at recording sites is lost due to partial RNA capture, recording site resection, or transcriptional silencing [10, 17, 18, 5, 6]. Finally, a mis-tuned “Cas9 editing rate” - the rate at which Cas9 induces heritable mutations used for lineage tracing - can lead to scenarios where there is a lack of mutation information sufficient for discerning relationships between cells. If the editing rate is too low, then “mutation-less edges” will occur in which a cell divides before it acquires a mutation, generating a irresolvable polytomy on the underlying phylogeny. If the editing rate is too high, then “mutation saturation” occurs in which the recording sites all acquire mutations before cell division ceases, making differentiation of the bottom of the phylogeny impossible [18]. As these complications are functions of parameters of the CRISPR-Cas9 lineage tracing system, a question of experimental design arises. Specifically, which configurations of the many experimental parameters involved in the CRISPR-Cas9 lineage tracing technology can alleviate these complications and make the problem of exact reconstruction more feasible? Our goal in this study is to address this question and identify configurations that are sufficient to theoretically guarantee exact reconstruction while also providing the accompanying phylogeny inference algorithms.

**Figure 1:**
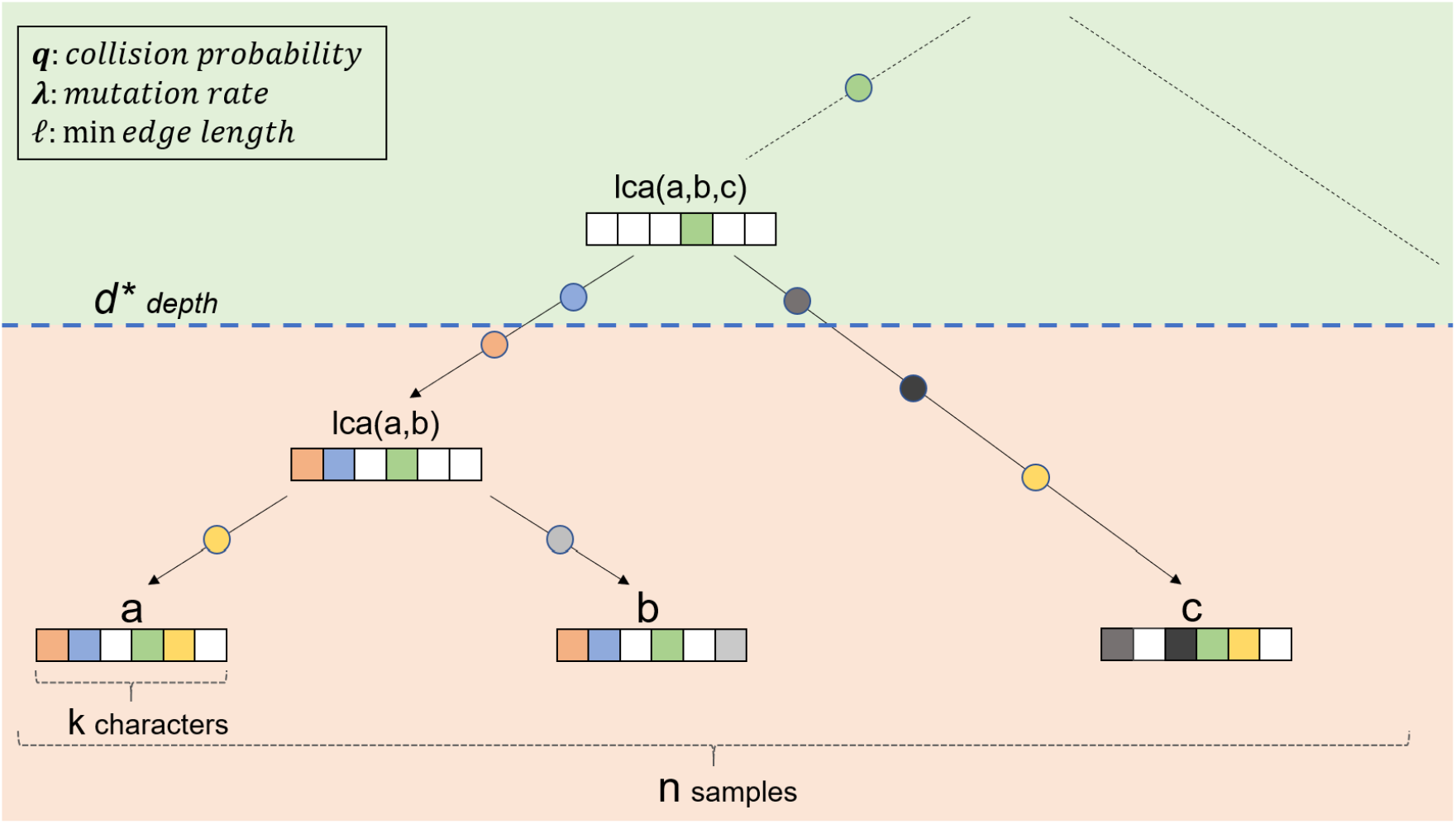
General problem setup. In particular, we note that *a* and *b* are the ingroup, whereas c is the outgroup. In addition, we point out some of the key variables used throughout this manuscript: *d*\*, *k*, *n*, *q*, *λ*, *ℓ*.

While there is extensive work establishing theoretical guarantees for exact reconstruction in other (arguably simpler) phylogenetic models, these models do not capture the specific features of CRISPR-Cas9 lineage tracing. One such example is the Cavender-Farris-Neyman (CFN) 2-state mdoel (a.k.a. binary Jukes-Cantor). In this setting, for *n* taxa, algorithms have been developed to exactly reconstruct subtrees with the number of characters 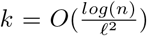 [19, 20, 21] if the length (duration) of every edge is greater than some value *ℓ*, and complete reconstruction is possible if the length of each edge is also upper-bounded by 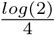 [22, 23]. Unfortunately, the CFN model cannot be readily applied to data of this type as it differs from the CRISPR-Cas9 settings in several critical ways: there are only two states (mutated, non-mutated) as opposed to the arbitrary state space of indels caused by Cas9 repair, reversibility of mutations is allowed as opposed to the irreversible mutations of Cas9, and there is no missing data [24, 25]. This motivates the development of theoretical bounds for the CRISPR-Cas9 lineage tracing model in particular.

Despite this need, to our knowledge there has yet to be any work directly exploring guarantees for exact reconstruction in the general CRISPR-Cas9 lineage tracing model. There do exist a few regimes where theoretical guarantees exist (e.g., under perfect phylogeny where every mutation occurs exactly once [26, 10]), but these regimes rarely exist in experimental settings. When these conditions are not met, other methods rely on criteria defined over the reconstructed phylogeny to guide reconstruction. Maximum-likelihood [27, 17], parsimony-based [28, 3, 10, 8] and distance-based [29, 30] methods optimize over likelihood, the minimum number of mutations, and variations of the ME (minimal evolution) criterion respectively [31, 32]. Unfortunately, optimizing these criteria does not necessarily correspond to reconstructing the correct tree. While there have been results regarding exact reconstruction for Neighbor-Joining in particular given certain error bounds between the true and observed distance metric [33, 34], there has yet to be work characterizing how and when mutation-based distances meet these criteria. Additionally, while existing studies have used simulations to provide insight into the relationship between reconstruction accuracy and experimental parameters [18, 10, 35], there has been limited theoretical exploration into CRISPR-Cas9 lineage tracing experimental design and how to optimize parameters in order to achieve accurate reconstruction.

In this paper, we derive such bounds for the CRISPR-Cas9 lineage tracing model. We develop two algorithms with theoretical guarantees for exact reconstruction of the underlying phylogenetic tree of a group of cells, showing that exact reconstruction can indeed be achieved with high probability given sufficient information capacity in the experimental parameters. In particular, we begin with an algorithm for a lower bounded edge constraint model whereby we prove that, in the absence of missing data, high probability of exact reconstruction can be achieved in polynomial time with 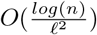 characters, matching the CFN 2-state model. The lower bound assumption translates to a reasonable assumption over the minimal time until cell division [36]. We further extend this algorithm and bound to account for missing data, showing that the same bounds still hold assuming a constant probability of missing data. We then consider the case of imposing an additional upper bound on edge lengths in our tree, to which we apply a bottom up approach that decreases the asymptotic number of characters required to 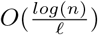 characters, improving on previous bounds. The upper bound corresponds to an assumption on the maximum time until cell division, which can be evaluated in lineage-traced populations, as they by definition should not be post-mitotic. Using these asymptotic bounds we characterize the dependence between the necessary number of characters and other experimental parameters, such as the mutation rate (controlled by guide affinity [9]), thus offering insight into how experimental design may be improved as the field develops. Lastly, we validate these relationships between *k* and the experimental parameters via a large set of simulations and present empirical bounds on the *k* required for exact reconstruction.

Taken together, our results provide a first theoretical analysis into the feasibility of phylogeny inference in the single-cell CRISPR-Cas9 based setting. As this field continues to grow and generate excitement as exemplified by the recent Allen Institute Dream Challenge [35], we anticipate that this work will inform the emerging dependencies between the experimental parameters and will serve to guide future studies both in terms of technology development (which parameters should be optimized) and tailoring the CRISPR-Cas9 lineage tracing system for specific case studies (e.g., dependent on the expected number of cell generations in the entire process). The algorithms and simulation engine presented here are available as an open source software in https://github.com/YosefLab/Cassiopeia.

## Problem Setup and Model Assumptions

In order to tackle the problems of guarantees on exact reconstruction and optimizing experimental design, it is helpful to consider a more abstract theoretical model. We begin with a single-cell with *k* unmutated characters, corresponding to *k* unedited recording sites at which CRISPR-Cas9 can induce mutations. This cell then undergoes cell division. Over time, characters may mutate from their unedited states at instantaneous rate *λ* according to an exponential distribution. While previous models have assumed a per-generation mutation rate [10, 37, 17], in our model, mutations occur independently from cell division. We believe this is a more accurate representation of current experimental regimes.

When a mutation occurs, the respective character adopts a state (corresponding to an indel) according to some probability distribution over the space of possible indels. We assume that once a character mutates, it can never change its state again and that this mutation will be inherited by all descendants of this cell. This irreversibility assumption is derived from the fact that after an indel is introduced at a recording site by Cas9, the guide RNA no longer has affinity at that site preventing future edits [18, 38]. After a set period of time has elapsed, a subset of the contemporary cells (leaves of the tree) are collected for sequencing. We denote the size of this set by *n*.

Finally, some proportion with expectation *p_d_* of each cell’s characters will have their states rendered indeterminable. This missing data may be due to low capture at the sequencing step and affect only one cell, or through resection (excision) or transcriptional silencing events which are inherited throughout the cell division process and persist in all descendant cells [10, 18, 5, 27, 6]. We refer to the former as stochastic missing data and the latter as heritable missing data.

Given the set of samples collected at the end of the experiment, the goal is to construct the phylogenetic tree that relates the observed cells to one another, based on their character/state information. Note that even though we collect a subset of the cells, the underlying “ground truth” phylogeny is still a binary tree. Formally, this problem can therefore be viewed as a character-based inference of a rooted phylogeny where the tree is binary, the root is unmutated, the mutations are irreversible, and some mutation data may be missing. In addition to offering algorithms to the problem, our goal is to shed light on the relationship between the various experimental parameters *k, q, n, λ*, and *p_d_* (summarized in Table 1), finding the regimes in which exact lineage reconstruction is tractable. We next explore the generative model in more detail.

**Table 1:** A summary of key model variables

| Parameters of Interest |  |
| --- | --- |
| Parameters | Description |
| $k$ | Number of characters |
| $n$ | Number of leaves in the tree. i.e. number of observable samples |
| $\ell$ | The minimum edge length on the ground truth phylogeny, normalized to the height of the tree |
| $\lambda$ | Rate at which each character mutates. Corresponds to rate at which recording sites are cut by Cas9 |
| $q$ | Probability of collision for independent mutation events |
| $p_d$ | The probability that the state at a given character/leaf pair is indeterminable because of missing data |
| $d^*$ | Depth such that any triplet rooted above $d^*$ that is well separated enough can be recovered with high probability. Normalized to the height of the tree |
| $\delta(d)$ | When all other parameters are fixed, $\delta(d)$ is a function that is proportional to the expected difference between ingroup-ingroup similarity and ingroup-outgroup similarity for triplet at depth $d$. It is the primary function used for the triplets oracle for making decision |

### Generative Process

Let 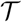 be a binary tree representing the ground truth phylogeny in a CRISPR-Cas9 lineage tracing experiment. Let *S* be the set of leaves in 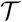, with each leaf representing a single cell in our input. Each edge (*u, v*) of 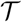 has a length, *l*(*u, v*), representing the duration of time between the two respective cell division events. Furthermore, we define the distance *dist*(*u, v*) between two vertices *u, v* as the sum of the lengths of the edges in the path between *u* and *v*. In addition, we denote the root node by *r* and say that a vertex *u* is at depth *d* if *dist*(*r, u*) = *d*. Finally we assume that the distance between the root and any leaf is equal to 1 (normalizing arbitrary time units), as all leaves are sampled at one time point, making the tree an ultrametric.

Each node in the tree has *k* independently evolving characters, each of which can take on states in {0, 1, *…m*}. Each character starts in an unedited state (0) at the root. Once a cell acquires a mutation at a certain character, this mutation is inherited by all descendants of that cell, and mutations cannot occur at that character in these descendants (irreversibility). For each character, the time it takes for a mutation to occur on a path in the tree is exponentially distributed with rate *λ*, and is independent of cell division events. That is, if *r* = 0^*k*^ is the root then the probability that a mutation occurs along the path from *r* to some downstream descendant vertex *u* for any particular character is

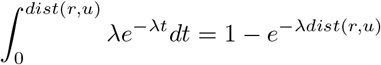

We assume that each character mutates independently of all other characters. Once a character mutates, it takes on state *j* ∈ {1, *…m*} with probability *q_j_*. Let 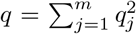 be the probability that two independent mutations at the same character index arrive at the same state. At the end of the experiment, each character in the leaves has a *p_d_* probability of becoming indeterminable and adopting the “missing” state. Finally, another measure of similarity between nodes, which we will use throughout, is derived from their mutation profiles. Here, we define by *s*(*u, v*) the number of mutations shared by nodes (cells) *u* and *v*. The definitions of the primary variables used in our analysis are summarized in Table 1 and Figure 1.

### Additional Definitions

For a triplet of nodes 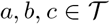, we use the notation of (*a, b|c*) to denote that *c* is the “outgroup”, namely that *LCA*(*a, b*) is a descendent of *LCA*(*a, b, c*), where *LCA* gives the lowest common ancestor in 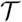. This leads us naturally to the concept of a triplets oracle.

#### Definition 1 (Triplets Oracle)

*We say that a function O*: *S*^3^ → *S is a triplets oracle if for every leaf triplet* (*a, b*|*c*) ∈ *S, O*(*a, b, c*) = *c.*

It is known that 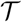 can be reconstructed exactly in *O*(*n* log *n*) time given a triplets oracle. Typically, it is unreasonable to expect to have exact triplet oracles in practical applications, so instead we define a relaxed version of this oracle.

#### Definition 2 ((*ℓ*\*, *d*\*) accurate partial oracle)

*We say that O is an* (*ℓ*\*, *d*\*) *accurate partial oracle, if for every triplet* (*a, b|c*) *such that depth*(*LCA*(*a, b, c*)) ≤ *d*\*, *then O*(*a, b, c*) *returns either c or Null. In addition, if it also follows that dist*(*LCA*(*a, b*), *LCA*(*a, b, c*)) ≥ *ℓ*\*, *then the oracle is guaranteed to return the correct answer, i.e. O*(*a, b, c*) = *c.*

In other words, the partial oracle does not return the wrong answer for triplets whose LCA is close to the root (max depth *d*\*). If in addition the triplet is separated by a distance of at least *ℓ*\* it is guaranteed to return the correct outgroup. In the remaining cases (e.g., triplets with an LCA far from the root, which are thus more difficult to resolve), the partial oracle can be wrong.

Throughout the paper, we will use the first order approximation 1 − *e*^−*x*^ ≈ *x* when suitable. We will also be using the following versions of Hoeffding’s inequality:

If *Y* ∼ *Bin*(*n, p*), and *μ* = *np* = *E*(*Y*), then we have:

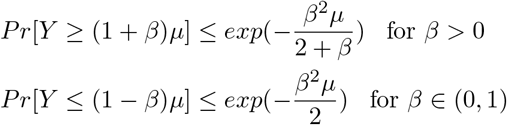

## Summary of Results

In the first section of this paper we show that in an experiment where each sample has sufficiently high number of characters and states (where the required number of characters and states depends on *λ, q, ℓ, d*\* and *p_d_*), that it is possible to construct (*ℓ, d*\*) partial oracles with high probability. Given these partial oracles, we have top-down algorithms that can exactly reconstruct 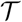 up to depth *d*\*, either with or without missing data. In particular, note that if *d*\* = 1 and *ℓ* is at most the minimum edge length, then an (*ℓ, d*\*) partial oracle is a triplets oracle, which leads to an exact reconstruction of 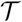 in *O*(*n* log *n*) time. In this paper, we study guarantees in full and partial exact reconstruction of the ground truth tree. We will give the guarantees about full reconstruction as corollaries of our main theorems as follows:

### Corollary 1

*Given a ground truth tree, 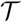, of height normalized to* 1 *under the lineage tracing model with k characters, n samples, minimum edge length ℓ, and constant mutation rate and state space, there exists a polynomial time algorithm to reconstruct* 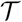 *with high probability when* 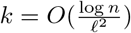.

Accounting for the possibility of missing data (incomplete information on the mutation profile of each cells in our study), we get:

### Corollary 2

*Under the same conditions above, assume that information of the state of any given character in a given cell can be masked with probability p_d_ independently for each character. In that case, there exists polynomial time algorithms to reconstruct* 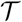 *with high probability when* 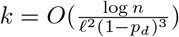.

As another extension, we consider a less constrained case with no lower bound on edge length. We show that in that case we can still get partial recovery as follows:

### Corollary 3

*Under the same conditions as in corollary 1 and with no lower bound on edge length ℓ, there exists a polynomial time algorithm that with high probability will return a tree which correctly resolves all triplets,* (*a, b*|*c*) *such that dist*(*LCA*(*a, b, c*), *LCA*(*a, b*) ≥ *ℓ*\* *when* 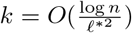.

Finally, we consider a more constrained case, where edge lengths are both upper- and lower-bounded. We demonstrate that it is possible to achieve a stronger lower bound on the number of characters required via a bottom-up algorithm. This part is based on an alternative strategy, in which we conduct a bottom up tree reconstruction without using an oracle. This alternative approach gives the following theoretical guarantee:

### Corollary 4

*If we assume that edge lengths are between ℓ and* 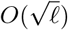 *then for a sufficiently low mutation rate, there exists polynomial time algorithms to reconstruct* 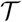 *with high probability when* 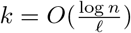.

## Partial Reconstruction of Phylogenies with Top-down Oracle-based Algorithms

For a triplet *a, b, c*, suppose WLOG that *s*(*a, b*) ≥ *s*(*b, c*) ≥ *s*(*a, c*). Our goal is to define a sufficiently high threshold *t* such if *s*(*a, b*) – max(*s*(*b, c*), *s*(*a, c*)) *> t* then with high probability, *c* is the outgroup. Since leaf node similarities can be readily computed from our input, this will help define a triplets oracle.

First, consider the case in which there is no missing data and all character states are determinable at the end of the experiment. Let (*a, b*|*c*) be a triplet where

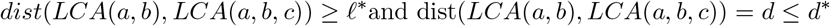

Given our assumptions on the mutation process, the similarities *s*(*a, c*) and *s*(*b, c*) have the same distribution, since *c* is the outgroup. Thus, we will focus on analyzing the quantity *E*[*s*(*a, b*) − *s*(*b, c*)] WLOG. Let *s_w_*(*u, v*) be the number of mutations shared by *u* and *v* that occurred after the point when the three lineages diverged from *LCA*(*u, v, w*). Then we have that *s*(*a, b*) – *s*(*b, c*) = *s_c_*(*a, b*) – *s_a_*(*b, c*) since any mutation that occurred before *LCA*(*a, b, c*) is inherited by all three nodes and will contribute equally to *s*(*a, b*) and *s*(*b, c*). The number of mutations shared by *a* and *b* after their divergence is distributed according to Binomial(*k, p*) where *p*, the probability of a given character having the same mutation in *a* and *b*, is

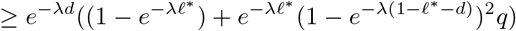

where *e*^−*λd*^ is the probability that the mutation did not occur before *LCA*(*a, b, c*). The left term inside the parentheses is the probability that the shared mutation came from a single event that occurred on the path from *LCA*(*a, b, c*) to *LCA*(*a, b*). The right term is the probability that the shared mutation came from two independent events that happened downstream of *LCA*(*a, b*) (i.e., homoplasy).

Considering there are *k* characters, we then have:

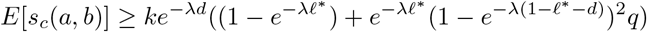

A similar computation shows that:

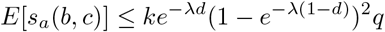

Thus, we have that:

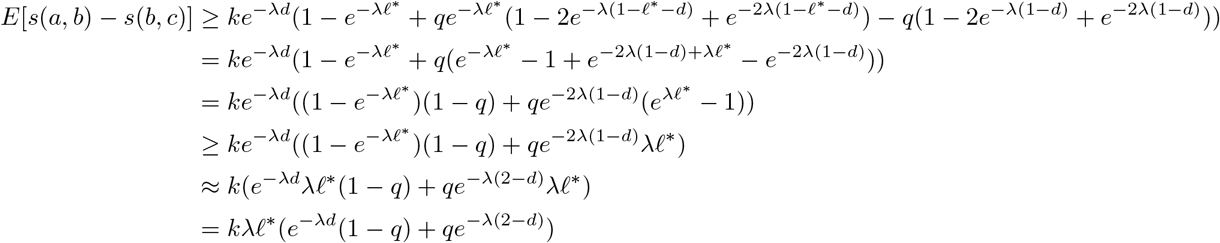

Let *δ*(*d*) = *e*^−*λd*^(1 − *q*) + *qe*^−*λ*(2−*d*)^. We then have that for any triplet (*a, b*|*c*), where *depth*(*LCA*(*a, b, c*)) = *d* and *dist*(*LCA*(*a, b, c*), *LCA*(*a, b*)) ≥ *ℓ*\*:

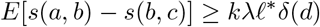

### Defining a (*ℓ*\*, *d*\*)−Oracle

We are now ready to define the decision rule that will be used by the partial oracle. Let *d*\* ∈ [0, 1] be an arbitrary depth to which we expect the oracle to be correct, and let *δ*\* = min_*x*∈[0*,d*\*]_ *δ*(*x*). Notably, the *δ*\* function has a closed form that depends on *λ, q*, and *d*\* (see appendix). For a particular triplet *a, b, c*, the oracle proceeds as follows:

i. Set a threshold 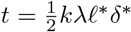.
ii. If there exists a pair *a, b* out of the triplet, such that *s*(*a, b*) – max(*s*(*a, c*), *s*(*b, c*)) *> t*, then return *c* as the outgroup. Otherwise return *Null*.

In the following we will prove that for a sufficiently large *k*, the function defined above is a (*ℓ*\*, *d*\*)–Oracle. In particular, let (*a, b|c*) be any triplet where *LCA*(*a, b, c*) is at depth *d < d*\*. We will prove that the following two conditions hold with high probability:

i. If *dist*(*LCA*(*a, b*), *LCA*(*a, b, c*)) ≥ *ℓ*\*, then *s*(*a, b*) − max(*s*(*b, c*), *s*(*a, c*)) *> t*
ii. In all cases, min(*s*(*b, c*), *s*(*a, c*)) − *s*(*a, b*) *< t*.

To see that conditions *i*) and *ii*) imply correctness of the (*ℓ*\*, *d*\*)-Oracle, note that the second condition guarantees that it is unlikely that the oracle will return the wrong answer when called on a triplet rooted at depth at most *d*\*. It will therefore return either the correct outgroup or *Null*. The first condition guarantees that if the triplet is also separated by a path of length at least *ℓ*\*, then the outgroup will be correctly returned.

Figure 5 in the appendix provides an empirical visualization of the (*ℓ*\*, *d*\*)-Oracle using simulations. The simulations mimic the CRISPR-Cas9 lineage tracing system, and are also described in the appendix.

Before computing the *k* necessary to make conditions *i*) and *ii*) hold, we first state the following lemma (proof in the appendix) which allow us to derive a worst case bound on the probability of triplets failing to satisfy condition *i*).

#### lemma 1

*Let* (*a, b|c*) *be a triplet with α* = *dist*(*LCA*(*a, b, c*), *LCA*(*a, b*))*. P* [*s*(*a, b*) – *s*(*b, c*) ≥ *t*] *is increasing with α.*

Now we can compute the *k* that ensures that both conditions *i*) and *ii*) are satisfied with high probability.

#### lemma 2

*Condition i*) *holds with probability at least* 1 − *ζ if we have the following guarantees on the parameters q, λ, ℓ*\*, *d*\* *and k*:

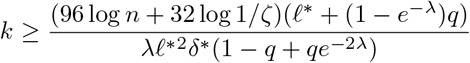

*Both conditions i and ii hold with probability at least* 1 − *ζ if we have:*

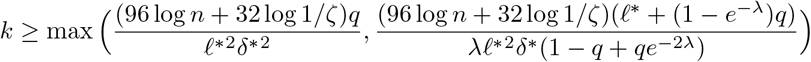

**Proof:** First, we will show that condition *i*) will hold with probability 1 − *ζ* if:

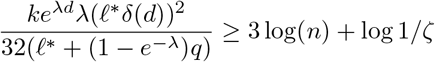

To see this, first note that *dist*(*LCA*(*a, b, c*), *LCA*(*b, c*)) ≥ *ℓ*\*. By lemma 1, we can WLOG assume that *dist*(*LCA*(*a, b, c*), *LCA*(*b, c*)) = *ℓ*\* because that is the worst case, i.e. the case where *P* [*s*(*a, b*) − *s*(*b, c*) ≥ *t*] is minimized. Any condition sufficient for this case will be sufficient overall. Let *Y* = *s_c_*(*a, b*) and *X* = *s_a_*(*b, c*). Since *E*(*Y*) − *E*(*X*) ≥ *kλℓ*\**δ*(*d*) ≥ *kλℓ*\**δ*\* = 2*t*, in order to ensure that *Y* − *X* ≥ *t*, it suffices to have *Y* − *X* ≥ *kλℓ*\**δ*(*d*)/2 which holds when *Y* ≥ *E*(*Y*) − *kλℓ*\**δ*(*d*)/4 and *X* ≤ *E*(*X*) + *kλℓ*\**δ*(*d*)/4. Thus, we have

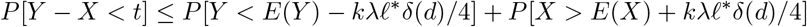

To bound the probability of both events using the above versions of Hoeffding’s inequality, we have:

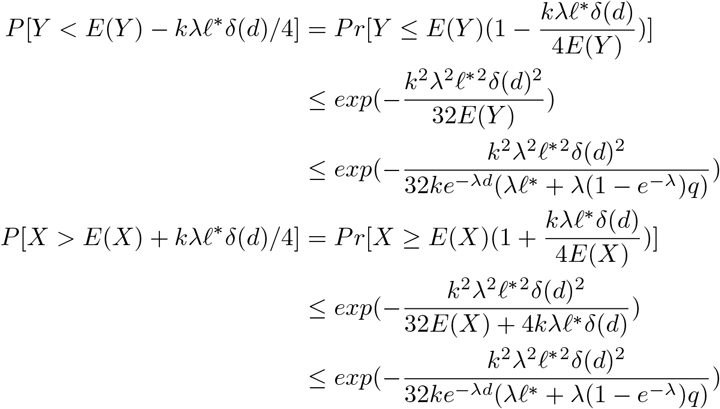

The last line follows from the fact that *δ*(*d*) ≤ *e*^−*λd*^ and *E*(*X*) ≤ *ke*^−*λd*^(1 − *e*^−*λ*^)^2^*q ≤ ke*^−*λd*^*λ*(1 − *e*^−*λ*^)*q*. Since *e^λd^δ*(*d*) = 1 − *q* + *qe*^−2*λ*(1−*d*)^ ≥ 1 − *q* + *qe*^−2*λ*^, in order to ensure that both bad events have probability at most *ζn*^−3^, it suffices to take

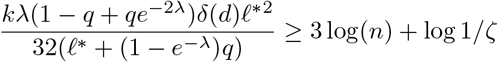

Since *δ*\* ≤ *δ*(*d*), the above condition holds as long as

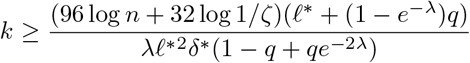

Applying the same argument to *s_c_*(*a, b*) − *s_b_*(*a, c*) and combining both results gives *P*[*s_c_*(*a*; *b*) – max(*s_b_*(*a*; *c*); *s_a_*(*b*; *c*)) < *t*] ≤ 4ζ*n*^−3^. Taking a union bound over all 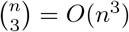 triplets, we see that the probability of condition *i*) failing for any triplet is at most ζ.

To get guarantees on the second condition, note that condition *i*) implies that condition *ii*) holds for all triplets separated by an edge of length at least *ℓ*\*. Thus we can focus on the triplets that are not covered by condition *i*). Let (*a, b|c*) be an arbitrary triplet such that *dist*(*LCA*(*a, b, c*), *LCA*(*a, b*)) *< ℓ*\* and again let *Y* = *s_c_*(*a, b*), *X* = *s_a_*(*b, c*) and *d* be the depth of *LCA*(*a, b, c*) (note that we are focusing WLOG on *s_a_*(*b, c*) since it has the same distribution as *s_b_*(*a, c*)). We want to show that with high probability, *X – Y < t*. Again, it suffices to upper bound *P* [*Y ≤ E*(*Y*) − *t/*2] and *P* [*X ≥ E*(*X*) − *t/*2] because *E*(*Y*) ≥ *E*(*X*). Note that we have already bounded the second quantity. To bound the first quantity, note that the worst case scenario is that *dist*(*LCA*(*a, b, c*), *LCA*(*a, b*)) is as small as possible. Since in this lemma we make no assumption on the minimal edge length, this quantity can be arbitrarily small and in the worst case, *dist*(*LCA*(*a, b, c*), *LCA*(*a, b*)) = 0, which means *Y* has the same distribution as *X*. Note that this case technically cannot happen as it would imply that 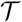 has a trifurcating branch but it is possible to get arbitrarily close to this case with no restrictions on edge lengths. This gives:

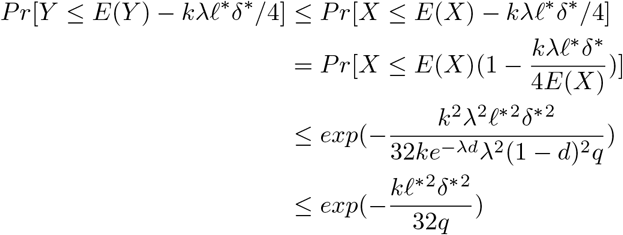

Thus, if we take:

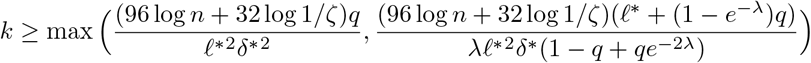

then we have *P*[*X* –*Y* ≥ *t*] < ζ*n*^−3^. By symmetry, this means *P*[*s_a_*(*b*; *c*) – *s_c_*(*a*; *b*) ≥ *t* ⋃ *s_b_*(*a*; *c*) – *s_c_*(*a*; *b*) ≥ *t*] ≤ 2*ζn*^−3^ Since we can union bound over one bad event of probability at most 2*ζn*^−3^ for each of the 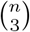 triplets, we have that conditions *i*) and *ii*) both hold with probability at least 1 − *ζ*.

An empirical demonstration and validation of the tightness of lemma 2 using simulations w.r.t. *λ* and *q* is provided in Figure 2, and w.r.t. *n* in appendix Figure 6. The simulations are described in the appendix.

**Figure 2:**
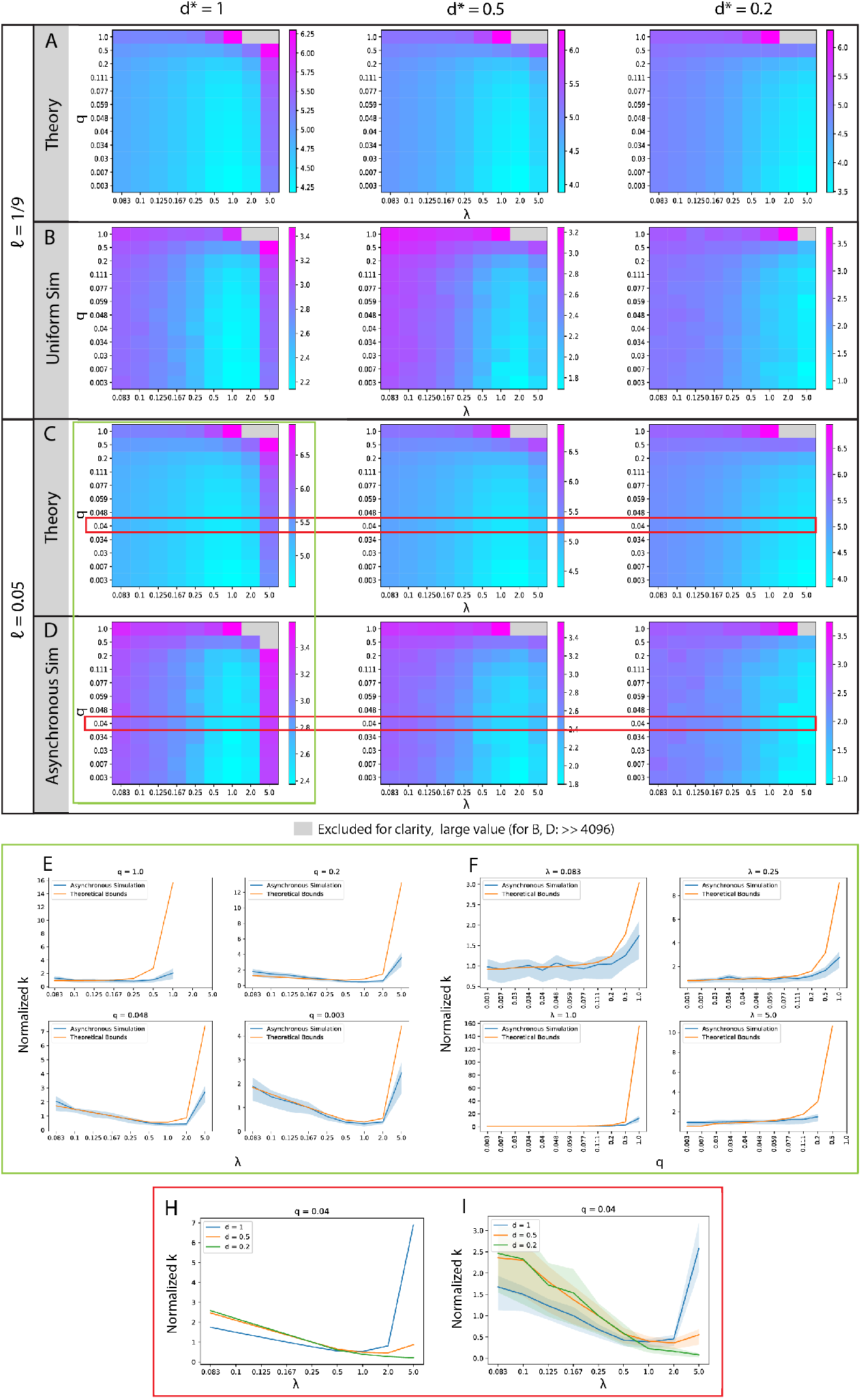
Comparing the Threshold Algorithm in theory and simulation. Simulated trees with 256 leaves, *n* = 256. (A, C) Theoretical sufficient lower bound on *k* required for 0.9 probability of perfect tree reconstruction up to depth *d* for varying values of *d*, *q* and *λ* for (A) *ℓ* = 1/9 and (C) *ℓ* = 0.05. (B, D) Minimum *k* required for 0.9 probability of perfect tree reconstruction in simulation with a cell division topology with (B) uniform edge lengths (*ℓ* = 1/9) and with (D) an asynchronous cell division topology (*ℓ* = 0.05). (A-D) Entries are *log*_10_ scaled. (E, F) Plots comparing the dependence of the minimum *k* in simulation with the theoretical bound on varying parameters (0.9 probability of perfect reconstruction, *ℓ* = 0.05, *d*\* = 1). We report the dependence of *k* on (E) *λ* for fixed values of *q* and (F) *q* for fixed values of *λ*. (H) Comparison of the dependence of the bound on *k* for 0.9 probability of perfect reconstruction on *λ* for various values of *d*. (I) Comparison of the dependence of minimum *k* for 0.9 probability of perfect reconstruction in the asynchronous simulation on *λ* for various values of *d*. (E, F, I) For ease of comparison, the values of *k* are rescaled by the median value of *k* in each line. (E, F, I) Point-wise 95% confidence intervals for the minimum *k* in simulation are generated from the regression coefficients using the delta method, see appendix.

### Sufficient Conditions for (*ℓ*\*, *d*\*)−Oracle in the Presence of Missing Data

#### General Regime

Now we consider the possibility of missing data (dropout) and give several simple strategies to handle it. In our analysis we consider two types of dropout events that may occur. A stochastic dropout is an event that occurs in and affects an individual cell (leaf), e.g., due to the limited sensitivity of single-cell RNA sequencing. A heritable dropout is an event that affects an entire clade, e.g., due to resection. Note that although we assume dropouts occur independently in each character, dropouts observed in the same character at two different cells are not necessarily independent as they could have originated from the same heritable dropout event. Now let *p_d_* be the probability that a particular character of a particular cell suffers either heritable or stochastic dropout. Let (*a, b|c*) be an arbitrary triplet and let *≤* be the probability that no dropout occurred in a particular character in either *a*, *b* or *c*. The probability that at least one cell of this triplet suffers a dropout at a particular character is maximized when the three cells share no common lineage and minimized when the three cells are phylogenetically proximal. We therefore have that (1 − *p_d_*)^3^ ≤ *ϵ* ≤ 1 − *p_d_* (see proof in the appendix). To account for this when revising our oracle definition, we now define *s*(*a, b*) as the number of characters shared by *a, b* that do not have dropout in either *a, b* or *c*. Note that in this case the definition of *s*(*a, b*) depends on *c* but is well defined for every triplet *a, b, c* so we can consider the same threshold-based triplet oracle before using this new similarity function. This means that for triplet (*a, b|c*), we have the following:

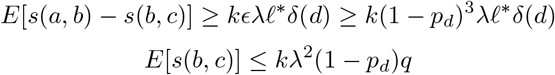

We can similarly define conditions *i*) and *ii*) with the threshold 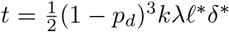, and see that if these conditions hold, then we have an (*ℓ*\*, *d*\*) partial oracle. We can also apply the same Chernoff bounds from the previous sections to get the following conditions on *k* to ensure conditions *i*) and *ii*) hold.

##### lemma 3

*(proof in the appendix) In the presence of missing data at a rate of p_d_, condition i*) *holds with probability at least* 1 − *ζ if we we have the following guarantees on the parameters q, λ, ℓ*\*, *d*\* *and k:*

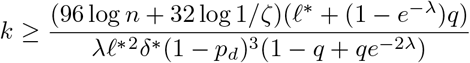

Both conditions *i*) and *ii*) hold with probability at least 1 − *ζ* if we have:

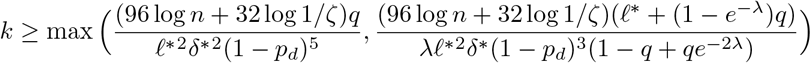

#### Alternative Strategy for Regime with only Stochastic Missing Data

Now we assume that missing data only occurs due to technical difficulties in reading out mutation sequences in cells, i.e. that only stochastic missing data occurs. Specifically, for any given cell and any given Cas9 recording site, we assume that the sequence of this site is missing from our data with probability *p_s_* and that this omission occurs independently from all other cells or mutation sites. With this definition, we consider a slightly modified oracle by defining *s*(*a, b*) as the number of mutations shared by *a* and *b* in characters that did not suffer dropout in either sample. Note that this definition of *s*(*a, b*) is independent of a third cell *c* unlike in the general case. Accordingly we revise conditions *i*) and *ii*) by setting the decision threshold *t* to be *k*(1 − *p_s_*)^2^*λℓδ*\*. These modified definitions lead to a more relaxed dependency between the mutation rate and the extent of missing data:

##### lemma 4

*(proof in the appendix): In the presence of stochastic missing data at a rate of p_s_, condition i*) *holds with probability at least* 1 − *ζ if we we have the following guarantees on the parameters q, λ, ℓ*\*, *d*\*, *and k:*

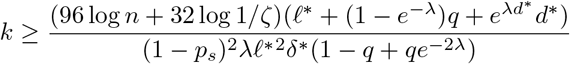

Both conditions *i* and *ii* hold with probability at least 1 − *ζ* if we have:

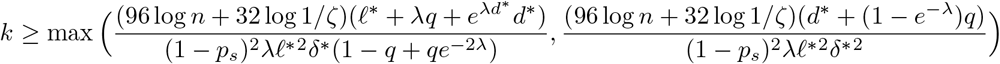

An empirical demonstration and validation of the tightness of lemma 4 using simulations is provided in appendix Figure 7.

## Top-down Algorithms for Oracle-Based Partial Tree Inference

Given our results on the correctness of the (*ℓ*\*, *d*\*)–Oracle, we are now ready to define the respective algorithm. Assuming *ℓ* bounded edge lengths in the ground truth tree, we use an oracle in which *ℓ*\* is set to *ℓ*. With that (*ℓ, d*\*)–Oracle, the algorithm guarantees accurate reconstruction up to depth *d*\* when given a sufficiently large *k*.

### Theorem 1

*In the lineage tracing regime, if*

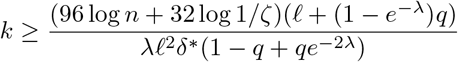

*and all edges in* 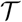 *at depth at most d*\* *have length at least ℓ, there exist polynomial time algorithms that return a tree which correctly resolves all triplets whose LCA is at depth at most d*\* *with probability at least* 1 − *ζ.*

**Proof:** By taking *ℓ*\* = *ℓ*, condition *i*) will hold for all triplets (*a, b|c*) whose LCAs are at depth at most *d*\* on 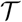 as *dist*(*LCA*(*a, b, c*), *LCA*(*a, b*)) ≥ *ℓ* for all triplets. The above bound on *k*, by lemma 2, implies condition *i*) is satisfied with probability 1 − *ζ*. It then suffices to show that whenever condition *i*) is satisfied, there exists a polynomial time algorithm that constructs a tree which correctly resolves all triplets whose LCAs are at depth at most *d*\*. We present a simple top-down recursive splitting algorithm which is guaranteed to return correct splits up to depth *d*\*. This algorithm has a runtime of *O*(*kn*^2^) for each recursive call, where *n* is the size of the input to the call.

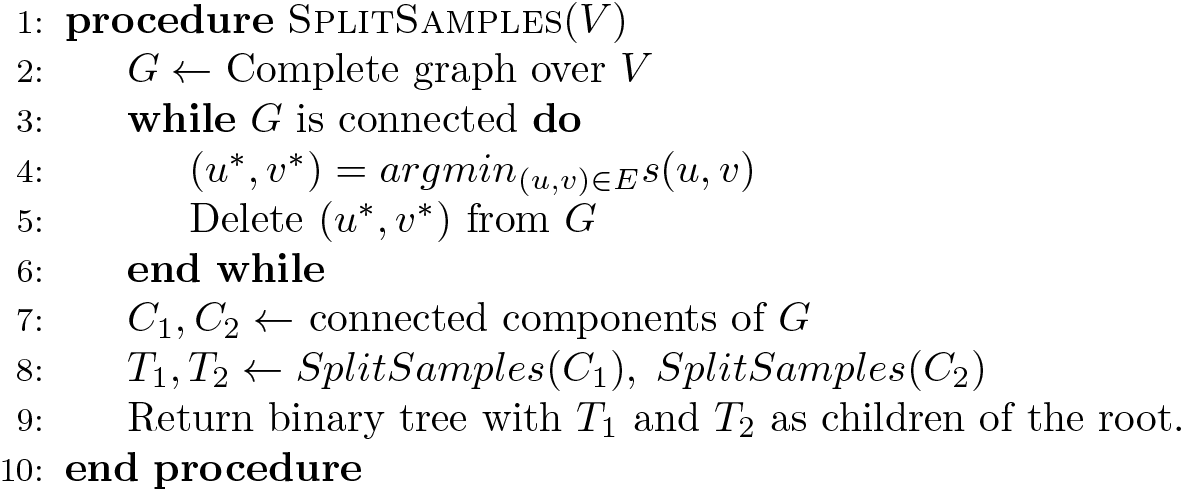
Threshold Algorithm

To prove correctness, let *V* be the set of samples at some recursive call in the algorithm. Let *r*′ be the LCA of *V* in 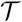 at depth *d ≤ d*\*. Let *L* and *R* be the samples in the left and right subtrees of *r*′ respectively. Let *a ∈ L* and *b ∈ R* be arbitrary. By condition *i*), we know that for any *c ∈ L*, *s*(*a, c*) *> s*(*a, b*) and for any *c*′ ∈ *R*, *s*(*b, c*′) *> s*(*a, b*). This means that whenever (*a, b*) is in the graph, and right after (*a, b*) is deleted from the graph if it ever happens, all neighbors of *a* and *b* will remain connected to them. Thus, the graph must remain connected until all all edges in the cut *L|R* are deleted, and *L* and *R* will still remain connected immediately after all cut edges are deleted, giving us the correct split. This means that the algorithm will keep returning correct splits as long as the LCA of all samples in a recursive call has depth at most *d*\*.

**Proof of Corollaries 1 and 2:** If we take *λ* and *q* as constants and *d*\* = 1, Theorem 1 implies that it is asymptotically sufficient to have 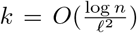 in order to ensure exact recovery of the entire ground truth tree with high probability. When dropouts are taken into account in the general case, the bound becomes 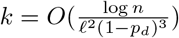 by lemma 3 because 1/(1 − *p*_*d*_)^3^ factor is needed to ensure condition *i*) still holds. Notably, when only stochastic dropouts are considered, the bound becomes 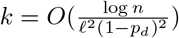 by lemma 4.

In the next theorem, we see that when we don’t have a lower bound on edge lengths, we can still construct a tree that correctly resolves all triplets that are well separated.

### Theorem 2

*In the lineage tracing regime, if the number of characters satisfy*

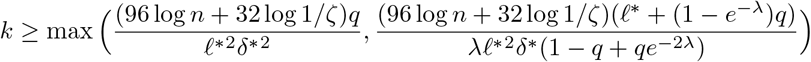

*and ℓ*\* *is an arbitrary parameter, then there exist polynomial time algorithms that return a tree which correctly resolves all triplets,* (*a, b*|*c*) *such that dist*(*LCA*(*a, b, c*), *LCA*(*a, b*) ≥ *ℓ*\* *and at depth*(*LCA*(*a, b, c*)) ≤ *d*\* *with probability at least* 1 − *ζ.*

**Proof:** Again, by lemma 2, it suffices to show that if conditions *i*) and *ii*) both hold, then there exists an algorithm which resolves all triplets, (*a, b|c*) such that *dist*(*LCA*(*a, b, c*), *LCA*(*a, b*) ≥ *ℓ*\* and at *depth*(*LCA*(*a, b, c*)) ≤ *d*\*. We can apply the classical Aho’s algorithm to recover a tree that is consistent with all triplets resolved by the (*ℓ*\*, *d*\*)–Oracle, which is guaranteed to us by conditions *i*) and *ii*). The algorithm is specified below for completeness; other supertree algorithms can be used here as well.

Let 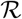 be the set of triplets for which we received a non-NULL answer from the oracle, and let *V* be the set of leaf nodes. Note that 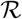 must include all triplets that are at depth at most *d*\* and whose internal nodes are separated by an path of length at least *ℓ*\*, since they satisfy condition *i*). It may also include incorrect triplets that are of depth more than *d*\*.

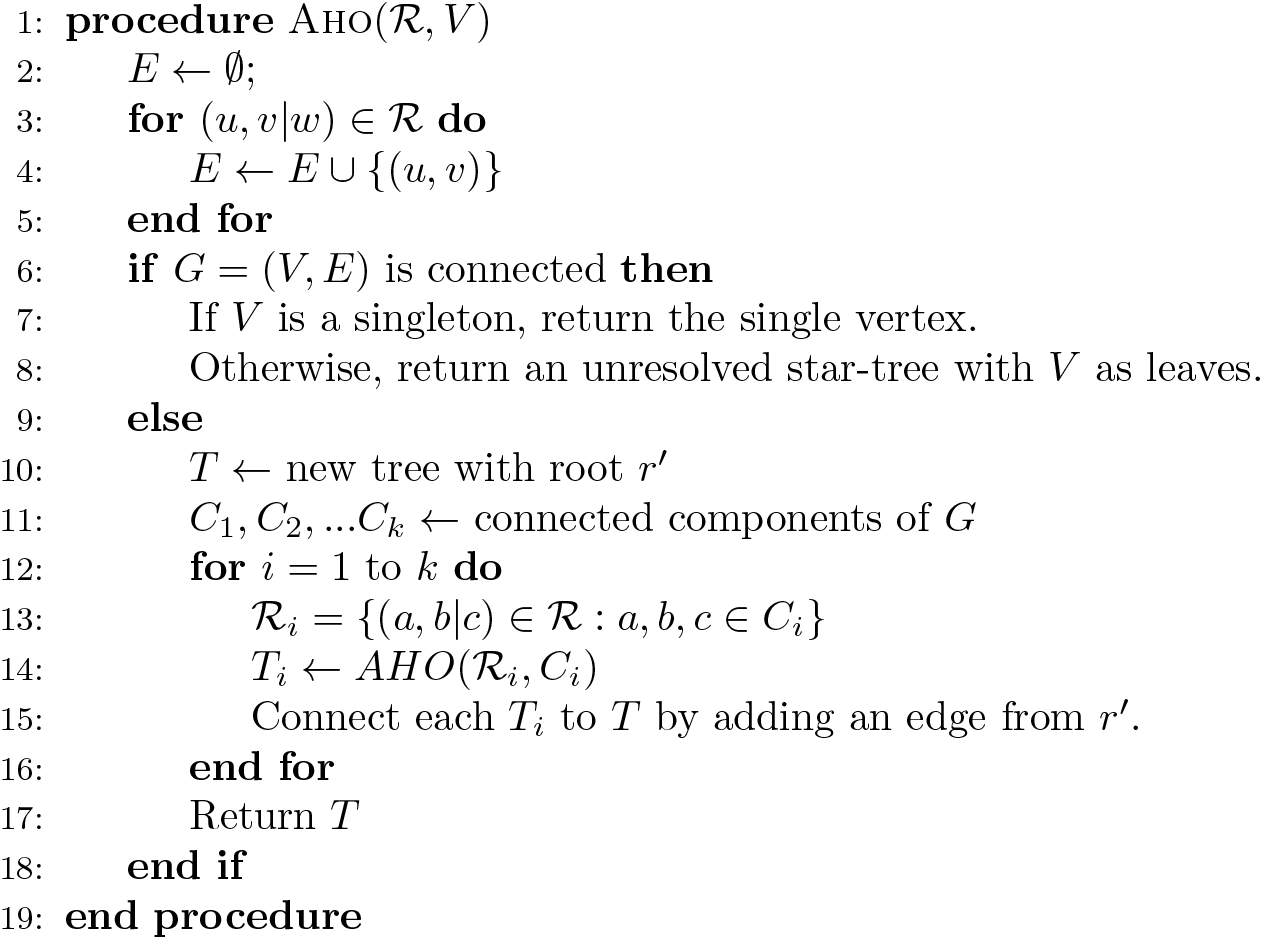
Aho’s Algorithm

To prove the correctness of the algorithm, we must show that if (*a, b|c*) is a triplet in the ground truth tree with *depth*(*LCA*(*a, b, c*)) ≤ *d*\* and *dist*(*LCA*(*a, b*), *LCA*(*a, b, c*)) ≥ *ℓ*\*, then (*a, b|c*) is correctly resolved in the tree returned by the algorithm. First, assume by contradiction that the triplet is represented wrongly (WLOG) as (*a, c|b*) in the returned tree. The presence of a wrong triplet (*a, c|b*) in the returned tree means that at some point, there was a recursive call on a set of leaves, *V* ∋ *a, b, c* such that *a, c* were in a connected component of *G* not containing *b*. However, condition *i*), combined with the assumption that *dist*(*LCA*(*a, b*), *LCA*(*a, b, c*)) ≥ *ℓ*\*, implies that if *a, b, c* ∈ *V* then there is an edge between *a* and *b*, which means *b* must be in the same connected component as *a* and *c*.

Next, assume by contradiction that *a, b, c* all have the same parent in the tree returned by the algorithm. First note that this cannot happen in line 15. This follows trivially since by definition *a* and *b* are initially in the same connected component. Therefore, the only way a trifurcation can happen is if a connected component that contains *a, b* and *c* is split into three or more component (with *a, b* and *c* on different components), each sent to a separate recursive call (line 14). This cannot happen since the presence of *a, b* and *c* entails the inclusion of an edge between *a* and *b* in that component (per line 4). This means that there was a recursive call on a set of leaves *V* ∋ *a, b, c* such that the connectivity graph, *G* over *V* is connected (i.e. the algorithm reached step 8). Let *r*′ be the LCA of all vertices of *V* in 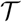. Let *L* and *R* be the vertices in *V* descended from the left and right children of *r*′ respectively. Since *r*′ *< d*\* condition *ii*) implies that any triplet with all leaves in *V* will be either correctly classified or assigned “Null” by the oracle. But, there are no edges between *L* and *R*, which means *V* is not connected, thus arriving at a contradiction once again. Thus, the only possibility is for (*a, b*|*c*) to be correctly classified in the inferred tree.

**Proof of corollary 3:** We take *λ* and *q* as constants and *d*\* = 1. Additionally, we make no lower bound assumptions on the edge length *ℓ* and take *ℓ*\* to be an arbitrary parameter. Theorem 2 then implies that it is asymptotically sufficient to have 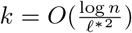 in order to ensure exact recovery of all triplets (*a, b*|*c*) such that *dist*(*LCA*(*a, b, c*), *LCA*(*a, b*) ≥ *ℓ*\* with high probability.

### Simulations for the Threshold Algorithm

Theorem 1 gives a lower bound on the number of characters *k* sufficient for exact phylogenetic reconstruction in the case where there is a minimum edge length *ℓ*. In order to both validate our asymptotic relationships between experimental parameters, as well as get a better sense for the number of characters that may be necessary in practice (a number that may be lower than our estimated of sufficiency), we turn to simulations. Specifically we chose to explore the empirical number of *k* necessary for exact reconstruction (all triplets resolved correctly) up to some depth *d*\* and as a function of the state collision probability *q* and the mutation rate *λ* (see Appendix for description of our simulations).

Figures 2A,C depict the dependence of *k* for high probability (0.9) of exact reconstruction with varying *λ* and *q* for 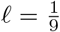 or 0.05. We observe that in the regions where *q > ℓ/λ*, the sufficient *k* increases sharply. This is since for lower values of *q* the asymptotic requirement for *k* becomes 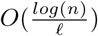, whereas for higher values of *q* we get the general result of 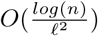. Additionally, we observe interesting behaviors in *λ*. In particular when *d*\* is large enough (requiring exact reconstruction of the entire ground truth phylogeny or its top half), both excessively small and large values of *λ* lead to a larger requirement for *k*. Intuitively this is due to the lack of mutations or due to mutation saturation, in both cases leading to less informative input (Figure 2H). When the goal becomes partial reconstruction of only the top (20%) of the phylogeny and *d*\* is small, *k* no longer increases with large *λ*. This is because *δ*\* in the denominator of the bound shifts from *e*^−*λ*^ to 1 as *d*\* → 0. Intuitively, towards the top of the phylogeny characters are yet to be saturated, allowing cells whose LCAs are near the top of the tree to be resolved. This suggests that saturation is less problematic if only distal relationships need to be resolved correctly, i.e. in the case of *d*\* *<<* 1.

In order to test the performance of the Threshold Algorithm in realistic settings, we simulate CRISPR-Cas9 induced phylogenies over two topological regimes: one with uniform edge lengths separating cell divisions with 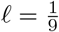 and one with an asynchronous cell division topology (*ℓ* = 0.05) described in the appendix (Figure 2B,D). The first regime aims to mimic a cell division process that has a regular molecular clock with bounded edges lengths both from above and below. The second regime is meant to mimic a more general stochastic cell division process that only has a minimum bound on edge length.

We find that the theoretical analysis and the simulations are consistent, both in terms of the direction dependence on the parameters, and on the inflection points at which the minimal *k* increases more rapidly. The largest discrepancies in the trends occur in the regions in which *λq* is high. In these regions the empirical increase in *k* is not nearly as sharp as the theoretical bound suggests. Hence the theoretical bound overestimates *k* relative to other values in these regions. Additionally, we observe that as *d*\* decreases the empirical *k* decreases (Figure 2I). This last result validates the trend in the bounds regarding *d*\*. Ultimately, the theoretical estimate predicts the empirical trends well, however, we do find that the absolute number of necessary characters (as found by the simulation) is much lower than the theoretical estimate.

Overall, these simulations provide validation that the asymptotic trends on *k* given by the theoretical parameters apply in realistic scenarios under the Threshold Algorithm. In addition, we provide necessary conditions for the number of characters required for exact reconstruction via the Threshold Algorithm for a series of parameter regimes.

## Upper-Bounded Edge Lengths and Bottom-Up Approaches

In the previous sections, we saw that when *λ* and *q* are fixed to be a constant, then the number of characters needed for exact recovery with high probability is 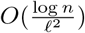, where *ℓ* is the minimum edge length. It can also be shown that if we are able to bound *λ* and *q* such that *λq ≤ ℓ* then the bound becomes tighter: 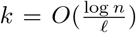. However, while *λ* can be controlled experimentally by calibrating the affinity of the lineage tracer’s guide RNAs, there is currently no way to control the entropy of the indel state distribution - a quantity which relies on the endogenous DNA repair process.

In order to achieve this tighter bound without direct dependence on *q*, we instead use an additional assumption on *ℓ* - namely that there is some maximal (in addition to a minimal) possible period of time between the birth of a given node and the birth of its parent. If our set of leaves includes all the cells in the phylogeny, then this translates to an upper bound on the time between cell divisions. In the more common scenario of sampling only a small subset of cells from a given clone, each edge in the ground truth tree can correspond to a series of cell division events. However, in either case, a strict upper bound on the length always exists, corresponding to the duration of the lineage tracing experiment. In the following section, we show that with such an upper bound on edge length, we can achieve exact recovery with high probability when 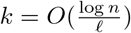, provided an upper bound on the probability of the most likely mutation (max_*j*_(*q_j_*)) and on *λq*. The latter bound can be less strict than *ℓ*, depending on the ratio between the lengths of the longest and shortest edges. More importantly, under these revised assumptions, we can achieve a bound on the minimal required *k* that is independent of the value of *q*.

### Theorem 3

*Let* 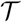 *be a tree of height* 1 *over n leaves. Let ℓ and c be constants such that each edge,* 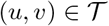 *has* 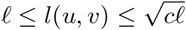. *Suppose that for each character we have*: 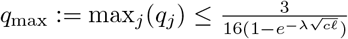 *and* 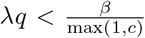 *where* 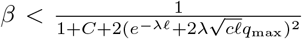 *and* 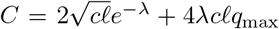. *Then there exists an algorithm that, with high probability, will recover* 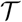 *if the number of characters satisfies:*

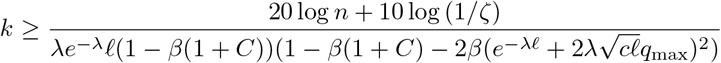

If we take the limit as *ℓ* → 0 or *n* → ∞ we get the following result:

### Corollary 5

*Let* 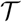 *be a tree of height* 1 *over a sufficiently large number of leaves n. Define ℓ, c and q*_max_ *as in Theorem 3 with similar bounds. If* 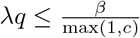 *where* 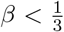 *then there exists an algorithm that can, with high probability, recover* 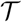 *if the number of characters satisfies:*

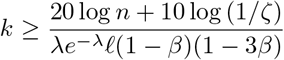

Note that the bound on *β* is not the tightest possible, and it was chosen to simplify calculations. Additionally, we present an alternative analysis in the appendix (Theorem 4) that yields a bound which has a looser constraint on the *λ* and *q* parameters as *ℓ* tends to 0.

**Proof of Theorem 3:** Consider the following greedy bottom-up algorithm which iteratively joins partially constructed subtrees by picking the pair with the most similar roots, and then joining them by inferring a new root by maximum parsimony. Let *S* denote the set of subtrees at any particular iteration. Let *T* ∈ *S* denote an inferred subtree, and let *r*(*T*) denote the root of that subtree. Let *r*(*T*)_*i*_ denote the state of the inferred *i^th^* character of *r*(*T*).

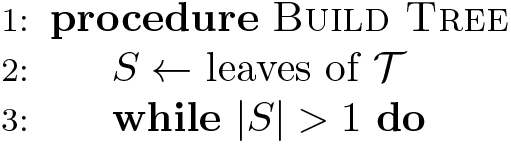
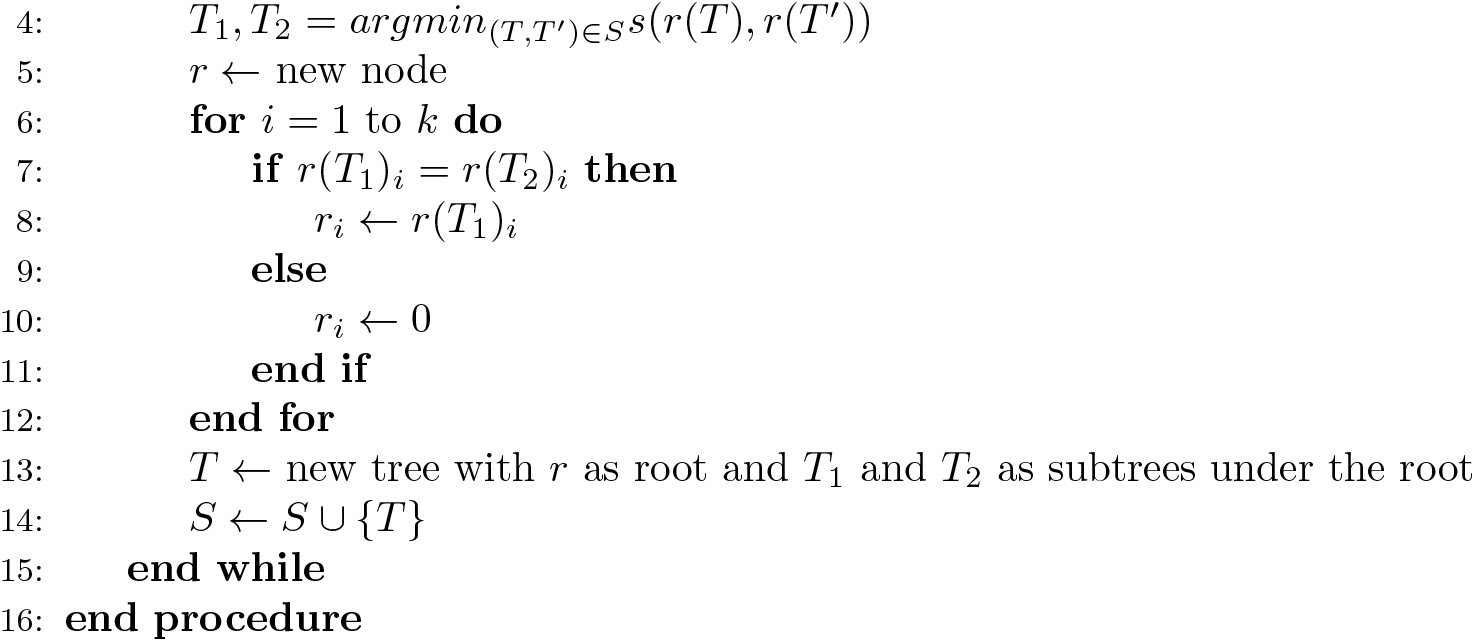
Bottom-Up Algorithm

(We note a similar algorithm has been presented in Sugino et. al [37], although no theoretical guarantees on accuracy are given in that work.)

In the following we show that under the conditions in Theorem 3, this algorithm correctly returns the ground truth tree 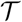 with high probability.

To show this, we must first bound the effects of incorrectly inferred mutations at internal nodes. Assume that *r*′ is an internal node generated in line 5 of the algorithm. Further assume that the tree rooted by *r*′ is correct (i.e., it is a sub-graph of 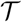). The process of assigning a mutation state to *r*′ (lines 6–12) may incorrectly include mutations that emerged independently at its two child trees. To account for this, we define *P_i,j_*(*r*′) to be the probability that every leaf beneath *r*′ has character *i* mutated to state *j*, given that character *i* is not mutated at *r*′. This probability is bounded according to the following lemma (proof in appendix).

### lemma 5

*Given the constraints on λ, ℓ and q in Theorem 3, it follows that* 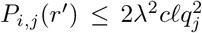 *for any internal node* 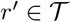, *character i and state j.*

To guarantee that the algorithm has a low probability of making mistakes in forming cherries (line 13), we focus on a pair of “active” nodes *u, w* in a given iteration of the algorithm (i.e., nodes that are roots of trees in the current set *S*; defined in lines 2 and 14). We assume that up until this point, the algorithm did not form any incorrect cherries (i.e., all trees in *S* are sub-graphs of 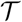). It suffices to show that with high probability, if *u* and *w* do not form a cherry in 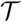, there must be some internal node, *v* on the path from *r*′ = *LCA*(*u, w*) to *u* such that *s*(*u, v*) *> s*(*u, w*) (note that *r*′ is LCA in 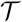). If this holds, then if *u, w* are the first incorrect pair to be merged, then there must be some descendent *u*′ of *v* such that *s*(*u, u*′) ≥ *s*(*u, v*) *> s*(*u, w*), contradicting the fact that *u, w* was the most similar pair (Figure 3). Note that *u* and *w* can be either leaves or internal nodes.

**Figure 3:**
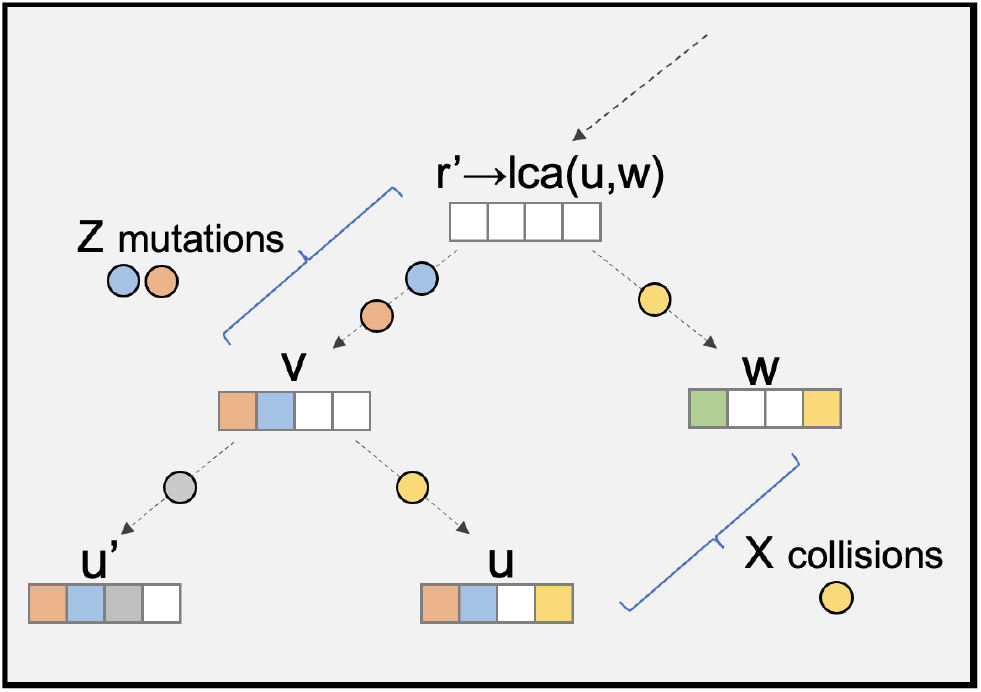
Primary variables for upper bounded special case. Let *u* and *w* be two arbitrary internal nodes that do not form a cherry. *v* is an ancestor of *u* that is a descendent of *r* = *LCA*(*u, w*). *Z* represents the number of mutations that occurred on the path from *r* to *v*. *X* represents the number of mutations that are shared between *u* and *w* due to homoplasy. If *Z > X*, then (*u, w*) will not be the first incorrect pair to be merged by the algorithm.

Let *d* be the depth of *r*′ and let *α* = *dist*(*r*′, *v*). Since the maximum edge length is 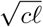, we have WLOG that 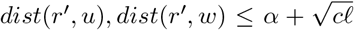 (if *dist*(*r*′, *w*) is greater than this quantity we can simply switch the roles of *u* and *w* and redefined *α* to be the distance from *r*′ to the parent of *w*). Let *Z* be the number of mutations that occurred on the path from *r*′ to *v*. This gives:

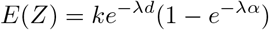

Note that due to irreversibility the mutations tallied in *Z* will be present in all nodes under *v*, including *u* and *u*′. Let *X* be the number of character assignments that are shared between *u* and *w* (per lines 7–11) and that are not present in *r*′ (per the ground truth). There are two ways in which these can happen: in the ground truth tree, a character can mutate to state *j* on the paths from *r*′ to *u* (or *w*), which occurs with probability at most 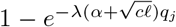. In that case, the mutation will be present in all downstream leaves, and thus assigned by the algorithm to *u* (or to *w*). If it did not mutate on that path, it could instead have mutated in enough places in the sub-tree rooted beneath *u* (or *w*) such that every leaf in that sub-tree has the mutation at that character. By lemma 5, we see that the probability of this occurring is at most 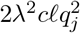. Requiring that in both *u* and *w* we have an occurrence of at lease one of these scenarios, we get:

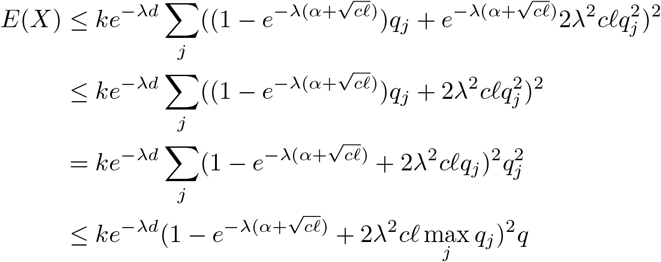

Let *q*_max_ = max_*j*_ *q*_*j*_. Assume *E*(*Z*) *> E*(*X*) and let Δ = *E*(*Z*) – *E*(*X*). Again by the above versions of Hoeffding’s inequality, we have the following concentration inequalities:

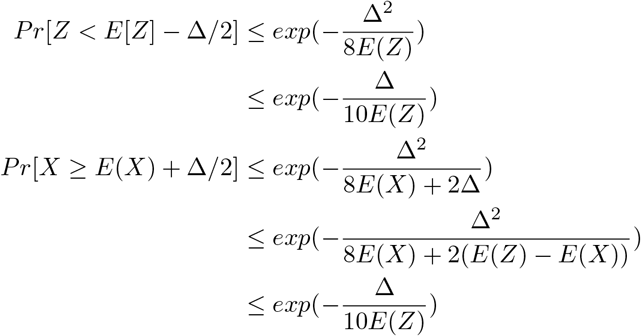

Suppose 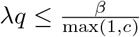. Let 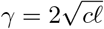. Then, we have:

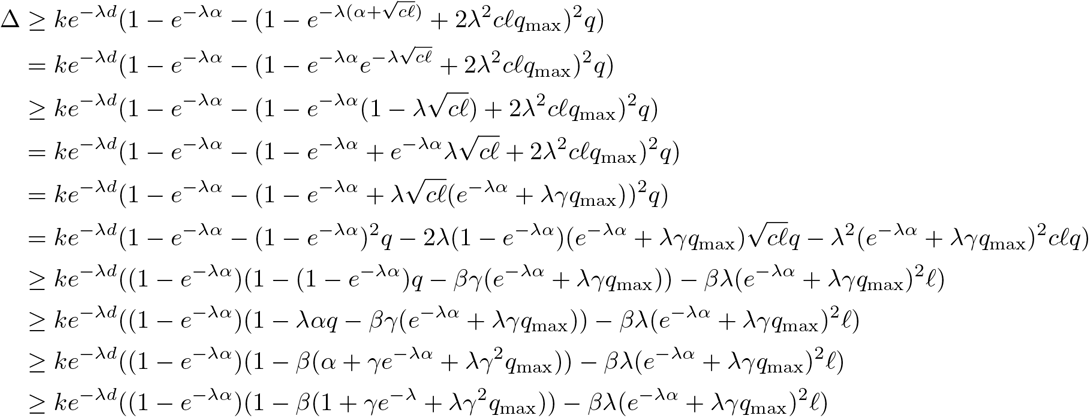

Where the last line follows from the fact that the maximum of the function *α* + *γe*^−*λα*^ occurs at *α* = 1.

Now, it remains to find a bound on *β* so that Δ *>* 0 and 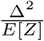 is lower bounded. Let *C* = *γe*^−*λ*^ + *λγ*^2^*q*_max_. To ensure that Δ *>* 0, we see that the last line from the previous block needs to be *>* 0. Taking this inequality and rearranging terms, it suffices to show that:

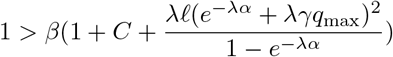

Note that

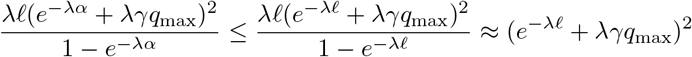

so our sufficient condition can be written as

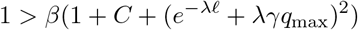

We lower bound 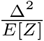 by the following:

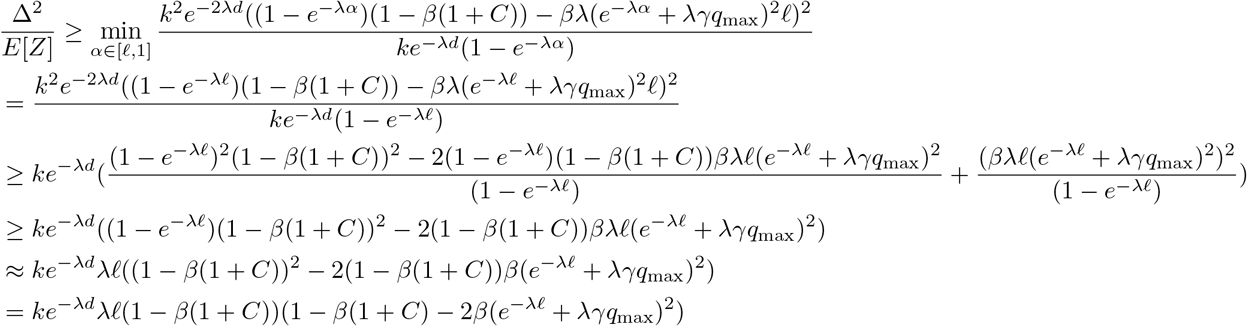

In order for this bound to be positive, we need

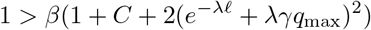

Note that this condition satisfies the above condition on *β* and thus would imply *E*[*Z*] *> E*[*X*]. Thus, if

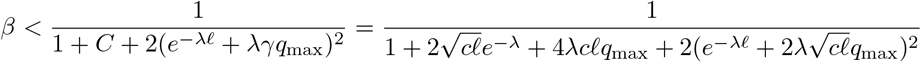

both Δ and 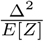 are strictly positive.

To bound the probability that *Z < E*[*Z*] − Δ/2 and *X* ≥ *E*(*X*) + Δ/2) is at most *n*^−2^*ζ*, we need:

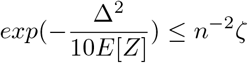

This is satisfied if *k* satisfies the following:

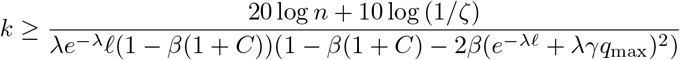

In other words, for any pair of vertices *u, w* that are not children of the same node, there will be a vertex *u*′ that such that *LCA*(*u, u*′) is a descendent of *LCA*(*u, w*) and *P* [*s*(*u, u*′) ≤ *s*(*u, w*)] ≤ 2*n*^−2^*ζ*. If *s*(*u, u*′) *> s*(*u, w*), then (*u, w*) cannot be the first pair of incorrectly joined vertices. Taking a union bound over at most *n*^2^/2 pairs of vertices, we see that with with probability at least 1 − *ζ*, there is no first pair of incorrectly joined vertices, which means the algorithm is correct.

An empirical demonstration and validation of the tightness of Theorem 3 using simulations w.r.t. *λ* and *q* is provided in Figure 4, and w.r.t. *n* in appendix Figure 6. The simulations are described in the appendix.

**Figure 4:**
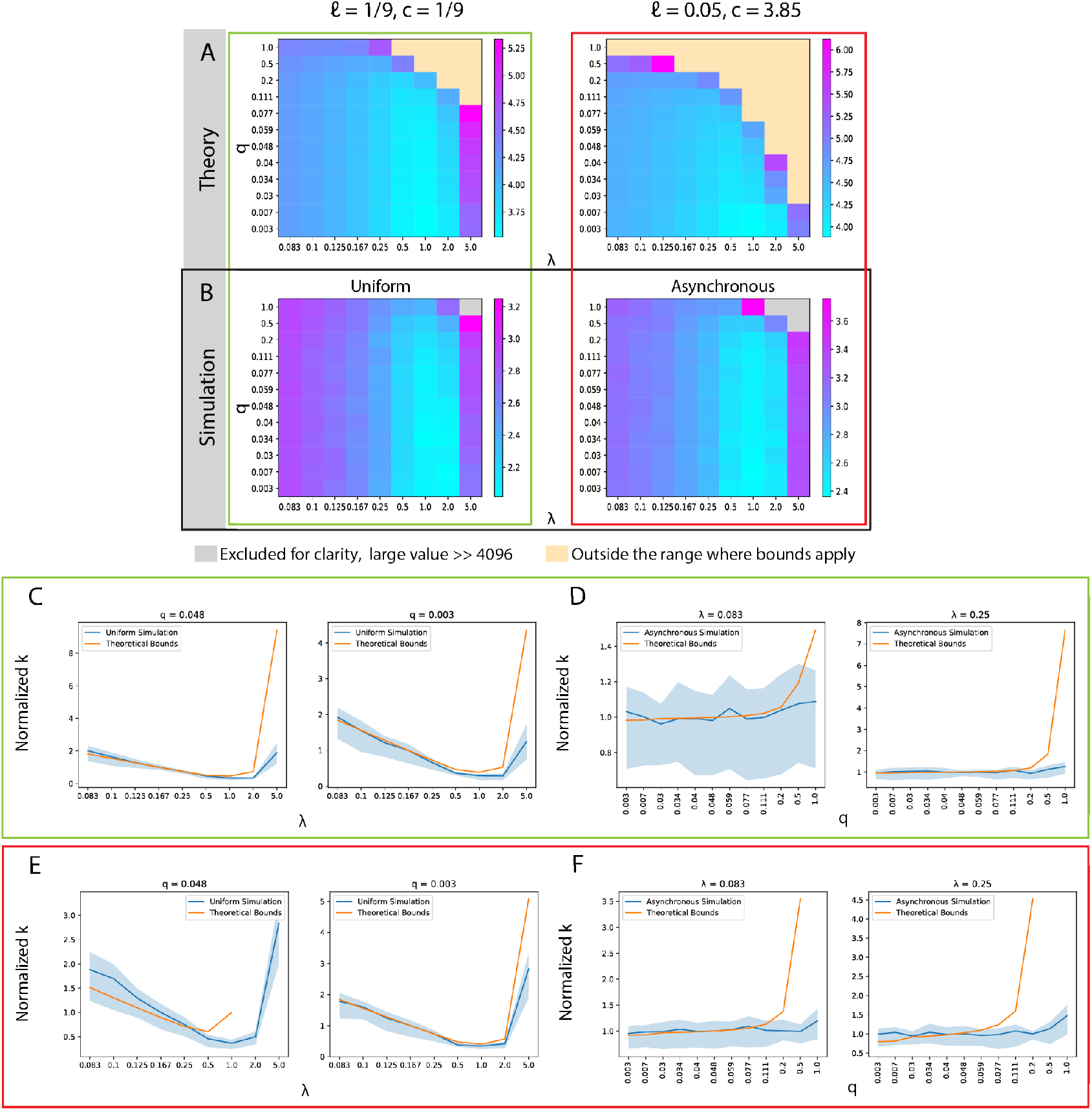
Comparing the Bottom-Up Algorithm in theory and simulation. Simulated trees with 256 leaves, *n* = 256. (A-B) Entries are *log*_10_ scaled. (A) Theoretical sufficient lower bound on *k* required for 0.9 probability of perfect tree reconstruction on varying values of *q* and *λ*, taking *β* = *λq* max(1, *c*). As the state distributions are uniform, *q_j_* = *q* for each value of *q*. Left: *ℓ* = 1/9 and *c* = 1/9 for comparison with the simulated case where the branch lengths are uniform. Right: *ℓ* = 0.05 and *c* ≈ 3.85 such that *>*99.99% of simulated branch lengths in the realistic simulation regime fall within the upper bound. (B) Minimum *k* required for 0.9 probability of perfect tree reconstruction in simulations, with Left: a cell division topology with uniform branch lengths, *ℓ* = 1/9 and Right: an asynchronous cell division topology (description in appendix), *ℓ* = 0.05. (C-F) Plots comparing the dependence of the minimum *k* in simulation with the theoretical lower bound on varying parameters (0.9 probability of perfect reconstruction). We report the dependence on (C, E) *λ* for fixed values of *q* and on (D, F) *q* for fixed values of *λ*, in simulations with uniform edge lengths (C-D) and with asynchronous topologies (E-F). (C-F) For ease of comparison, the values of *k* are rescaled by the median value of *k* in each line. Point-wise 95% confidence intervals for the minimum *k* in simulation are generated from the regression coefficients using the delta method, see appendix.

**Figure 5:**
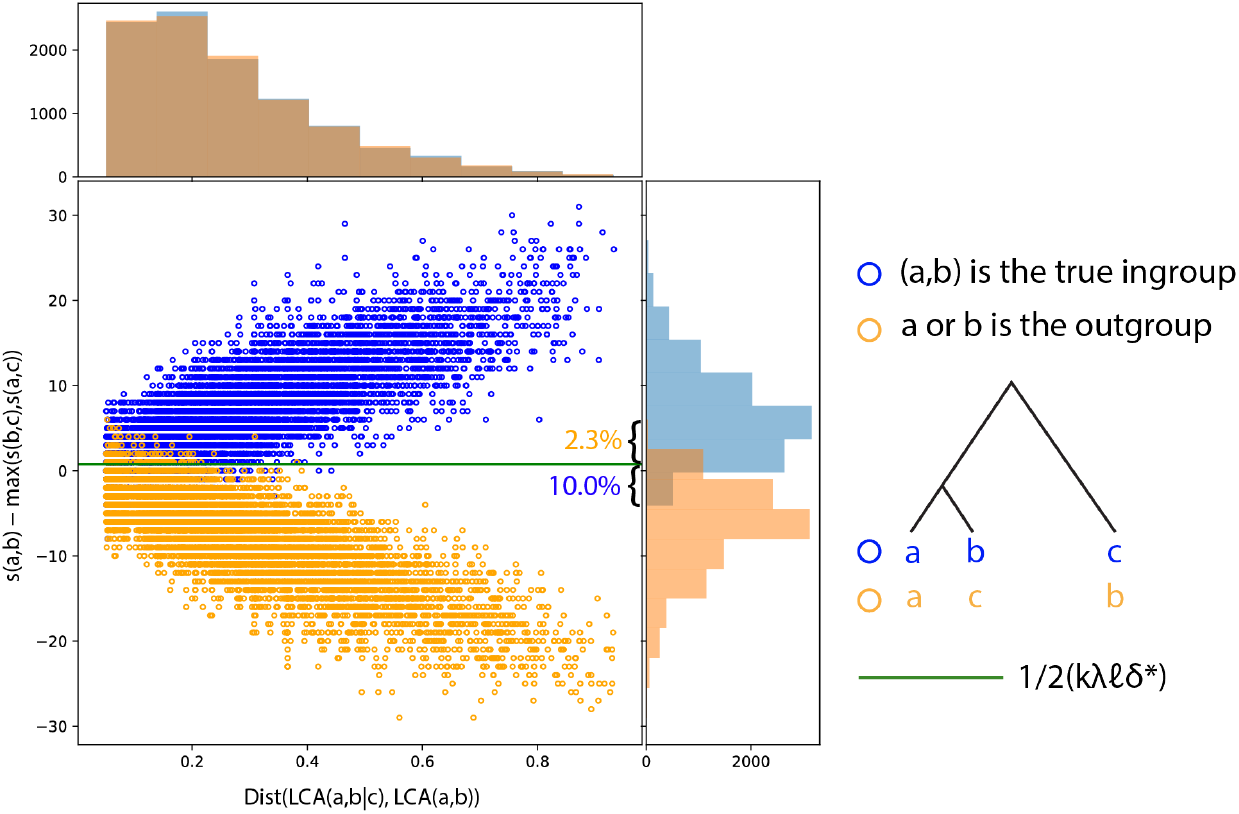
Visualizing the (*ℓ*\*, *d*\*)–Oracle decision rule in simulation. We sample 100 triplets uniformly for each of 100 trees simulated under the Asynchronous simulation framework described in the appendix with *k* = 10, *λ* = 0.5, *q* = 0.05 and *n* ≈ 256. For each triplet, we plot *s*(*a, b*) – max(*s*(*a, c*), *s*(*b, c*)) for two cases: when (*a, b*) notates the true ingroup of the triplet (blue) and 2) when (*a, b*) notates one member of the ingroup and the outgroup (orange). The histograms show the density of each case for each triplet along the axes.

**Figure 6:**
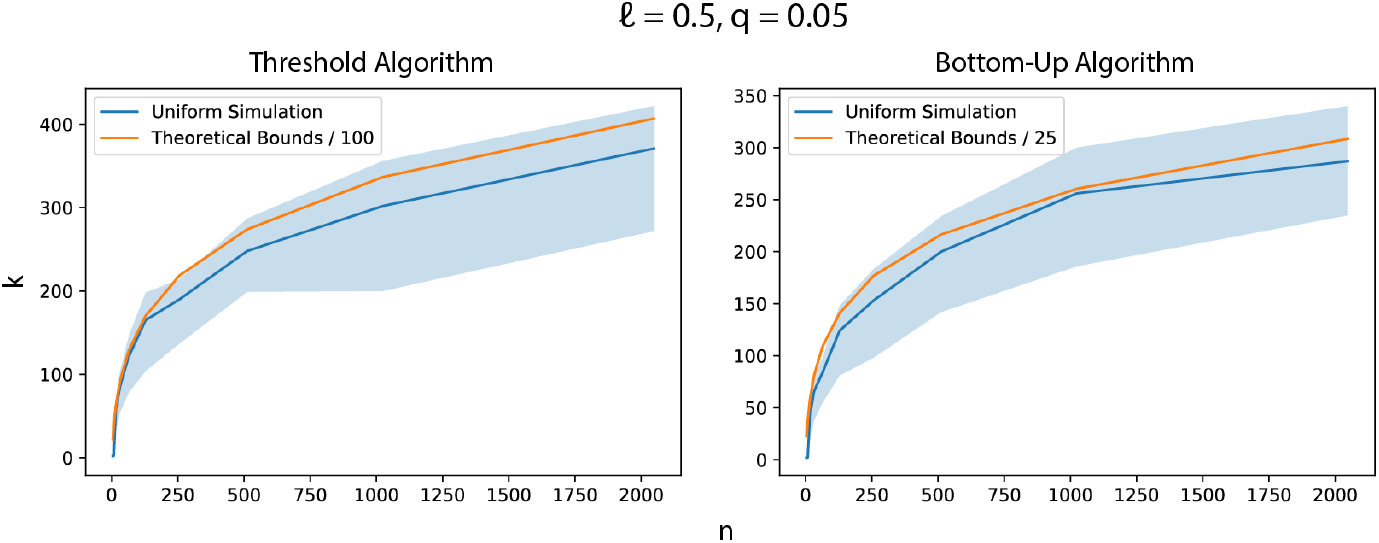
Comparing the dependence of minimum necessary *k* on *n* of the Threshold Algorithm (left) and the Bottom-Up Algorithm (right) with the theoretical bounds for each case (90% of full reconstruction). Simulations performed with the uniform edge length regime for trees of size 2^2^, 2^3^, *…,* 2^11^ leaves. For each value of *n*, edge lengths were re-scaled to be 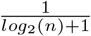 to maintain uniform edge lengths. The bounds are rescaled by a constant factor, 100 for the Threshold Algorithm and 25 for the Bottom-Up Algorithm. Point-wise 95% confidence intervals are generated from the regression coefficients using the delta method, see appendix.

**Figure 7:**
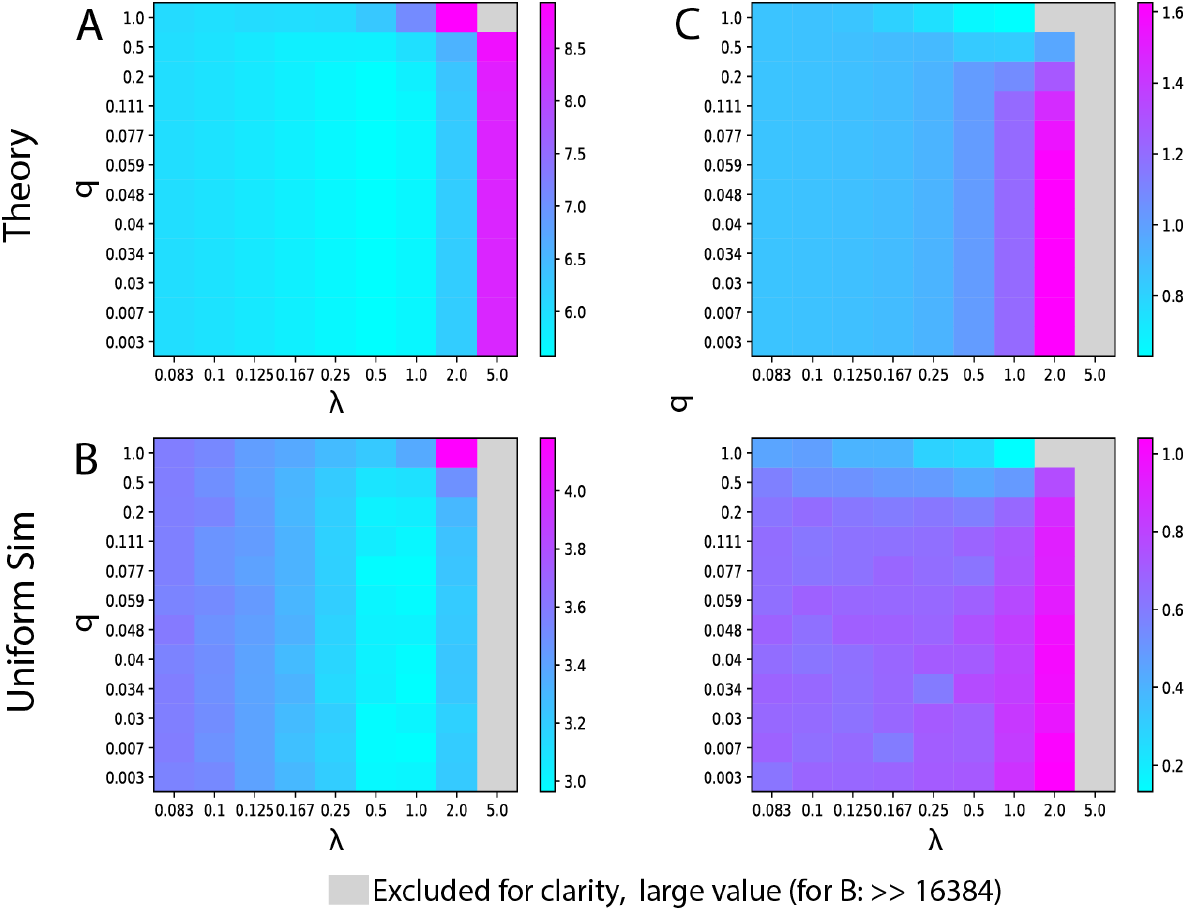
Comparison of the Threshold Algorithm in simulation and theory in the case of missing data, using the missing data strategy outlined in lemma 4. Simulated trees with 256 leaves, *n* = 256. Entries are *log*_10_ scaled. (A) Theoretical lower bound on *k* required for 0.9 probability of perfect tree reconstruction for varying values of *q* and *λ* in the case of missing data, with *ℓ* = 1/9, *d*\* = 1, and *p_s_* = 0.1. (B) Minimum *k* required for 0.9 probability of perfect reconstruction in simulation with a cell division topology with uniform branch lengths, *ℓ* = 1/9, *d*\* = 1, and *p_s_* = 0.1. (C) Scalar difference in the values of *k* in the case with and without missing data. Top: Difference in theoretical bounds for *k* for 0.9 probability of perfect reconstruction, *ℓ* = 1/9, *d*\* = 1, and *p_s_* = 0.1. Bottom: Difference in minimum *k* required for 0.9 probability of perfect reconstruction in simulation with a cell division topology with uniform branch lengths, *ℓ* = 1/9, *d*\* = 1, and *p_s_* = 0.1.

### Proof of Corollaries 4 and 5

The Bottom-Up Algorithm shows that there exists a polynomial time algorithm that ensures 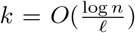 characters is sufficient asymptotically for exact recovery of the tree. As *n* → ∞ we get that *ℓ* → 0 and *C* → 0. With these we get that 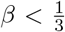. With the simpler bound on *β*, we also get that 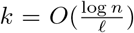 characters are sufficient asymptotically for exact recovery of the tree.

#### Simulations for the Bottom-Up Algorithm

Similar to the simulations for the Threshold Algorithm, we begin by Examining the theoretical bounds for the necessary *k*. In particular, Figure 4A visualizes the bound for *k* across varying values of *λ* and *q* for high probability (0.9) of exact reconstruction. We consider two regimes: one with *ℓ* = 1/9, *c* = 1/9 and one with *ℓ* = 0.05, *c* ≈ 3.85. Since *q* does not explicitly appear in the bound for *k*, we instead use it to define a value for *β*, using its lower bound: *β* := *λq · max*(1, *c*). Plugging this in provides a lower bound for the necessary *k*, which we plot here. Regions where the lower bound on *β* becomes larger than its upper-bound requirement (per theorem 3) are excluded.

From this figure it can be seen that *k* depends on *λ* in the same way as in Theorem 1. That is, *k* increases significantly for both excessively small and large values of *λ*. However, there is a contrast in the dependence of *k* on *q*. Although in the bound for Theorem 3 *k* does increase with *q* through the dependence of *β*, *k* is not as sensitive to large values of *q* as in Theorem 1. Further, as the bound is quadratic in 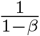 the *k* increases rapidly with respect to *β* := *λq* · *max*(1, *c*).

We tested the Bottom-Up Algorithm in the same simulation regimes (same tree and lineage tracing parameters) as the Threshold Algorithm (Figure 4B). Concordant with the theoretical results, we observed that the minimum required *k* is less sensitive to *q*, compared with the Threshold Algorithm. Furthermore, in both results we see similar trends in dependence on *λ* (Figure 4C, E), and *q* (Figure 4D, F). The main discrepancy between the theory and the simulation occurs where *β* := *λq · max*(1, *c*) approaches our upper bound for *β* (i.e., the values that border the regions that were excluded from Figure 4A). In those cases,we see that the theoretical bound is looser and overestimates *k* relative to the simulations.

Again, these simulations validate the relationships observed in the asymptotic trends on *k* and give tighter empirical conditions on the necessary *k* for exact reconstruction. In addition, we observe that the empirical necessary *k* in the Bottom-Up Algorithm is overall lower than that of the Threshold Algorithm, except the cases of non-uniform edge length with high value of *c* in which the minimal *k* is comparable (Figure 4A-B right, and Figure 4E-F). These results suggest that the Bottom-Up Algorithm can achieve exact reconstruction with fewer characters empirically than the Threshold Algorithm, but requires that the variance in the division times of the ground truth phylogeny to be small (corresponding to the assumption on upper bounded edge lengths).

## Discussion

In this paper we have established sufficient conditions for high probability of exact reconstruction of the ground truth phylogeny in the CRISPR-Cas9 lineage tracing setting. These guarantees show that despite complications with the lineage tracing process such as homoplasy, missing data, and lack of mutation information, exact reconstruction can still be achieved given sufficient information capacity in the experiment (as measured by the number of recording sites). In addition to showing the feasibility of exact reconstruction, these theoretical results relate the difficulty of the reconstruction problem in the number of sufficient characters to the experimental parameters. We anticipate these results can inform researchers as to how to reduce the number of necessary characters or best aid downstream reconstruction of the phylogeny given the available number of characters through careful engineering of CRISPR-Cas9 lineage tracing experiments.

The theoretical results shown here provide insight into how the CRISPR-Cas9 lineage tracing experimental parameters relate to the reconstruction problem. One key insight is that for exact reconstruction, a mutation rate that is neither too high or low gives the best results, which is in line with the intuition that a middling rate balances mutation saturation and mutation-less edges. We also formalize the intuition that having a state distribution with a low rate of collision *q* makes the reconstruction problem easier in avoiding homoplasy. Additionally we show the difficulty that missing data poses for the problem of exact reconstruction. Using the presented methods, the number of additional characters needed to overcome missing data is cubic (quadratic in the case of only stochastic missing data) in 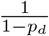, where *p_d_* is the probability of missing data. A final key result is that the (*ℓ, d*)-oracle in the Top-down Algorithms allows researchers to tailor the granularity of their reconstruction accuracy to what is achievable given the number of available characters. Here, substantially fewer characters are required if one is only interested in correctly resolving triplets that diverged early in the tree (small *d*\*) and well-separated triplets (large *ℓ*\*, regardless of the true minimum edge length *ℓ*).

Although having more characters is preferable, we recognize that currently there are practical limits on the number of recording sites that can be incorporated into CRISPR-Cas9 systems. Current methods to incorporate recording sites into the genome (such as lentiviral transduction [3, 10, 12] or transposition [5, 6, 9, 11, 14]) are limited by the low uptake of these sites into the progenitor cell, only offering on the scale of tens of recording sites [10, 6, 5, 12, 3, 9, 8, 7]. One alternate technology of particular interest is the base-editor, which uses a modified Cas9 complex to induce direct base-pair substitutions [39]. Base-editors, while yet to be explored in lineage tracing contexts, have the potential to offer one hundred or more editable sites [40], although careful engineering is required as *q* is high in these regimes owing to the limited state space of nucleotide outcomes. Ultimately though, we see in our simulations that even this increase in characters is far insufficient for exact reconstruction in most settings, especially considering the considerable amounts of missing data and the large number of samples (*n*) that we see in real CRISPR-Cas9 lineage tracing experiments. We thus challenge the field to develop systems that allow for considerably more characters.

The limitations in adding more characters motivates the optimization of the other experimental parameters in engineering CRISPR-Cas9 lineage tracing experiments. We discuss here current and potential strategies for engineering the discussed parameters: the editing rate, the collision probability, and the missing data probability. There is a large body of literature showing that the editing activity of Cas9 (as in lineage tracing experiments) can be tuned with relative precision using mismatches between the guide RNA and recording site [9, 18, 10, 12, 14]. In regards to the collision probability, experimenters are currently unable to dictate the collision rate in state outcomes due to the random indel outcomes of Cas9 editing. Recent strategies - such as pairing terminal deoxynucleotidyl transferase (TdT) with Cas9 [16] - have shown potential to increase indel diversity. Further, prime editors offer an avenue to more finely control the state distribution by dictating a-priori which indel will result in a given edit, though this technology has yet to be adopted for lineage tracing [41, 42, 43]. Fortunately, current CRISPR-Cas9 lineage tracing systems are capable of generating “un-problematic indel distributions”. In those cases, the collision probabilities *q* lie outside of the range where *k* explodes with *q* [9, 10, 12]. Additionally, we see in the bounds and simulations that decreasing *q* has diminishing returns on *k*. Taken together, in designing CRISPR-Cas9 lineage tracing systems effort is better put on carefullY ≤ Engineering the Cas-9 cutting rate than optimizing the state distribution.

Unfortunately, current strategies to control for missing data experimentally are more limited. Experimenters are at the mercy of the efficacy of single-cell assays in the case of low capture (leading to stochastic missing data), but can attempt to control the rate of resection and transcriptional silencing (both of which leading to heritable missing data). Recent designs have mostly relied on distributed designs to reduce the rate of resection, utilizing many “cassettes” (DNA segments that contain many proximal recording sites) with a small number of recording sites per cassette [8, 9, 10, 12]. Although not addressed in current designs, transcriptional silencing can be potentially limited by placing recording sites in regions of the genome that are more robust to silencing (“safe-harbor” regions) using emerging methods for guided transposition [44].

In addition to motivating the design of CRISPR-Cas9 lineage tracing experiments, our model motivates theoretical and algorithmic development for these systems. The sufficient bounds that we reach in our asymptotic analyses are not tight, as demonstrated by simulation. We believe that these bounds can likely be further improved to give a better sense of the necessary *k* analytically. Future approaches may take advantage of aspects of the model or engineering designs that are not leveraged in this work. For example, in our analysis we assume that the mutation rate *λ* is constant throughout the entire phylogeny and across recording sites. However, using a gradually increasing mutation rate or designing characters with variable rates, whose affinity is estimated a-priori may lead to better reconstruction results. Such a design can simultaneously alleviate issues of mutation saturation near the leaves of the tree as well as lack of sufficient mutations near the root. We also assume that the characters mutate and acquire missing data independently, although indels and missing data events can span multiple recording sites. Future approaches could take advantage of the structure present in these multi-site events. Finally, although our analysis handles missing data by ignoring missing characters, the structure of heritable data offers additional information that could be better leveraged (i.e., utilized in the same way as any other mutation). The challenge, naturally, is to distinguish between the two types of missing data.

Here, we perform a first theoretical analysis of CRISPR-Cas9 phylogenetic reconstruction. In doing so, we have developed a generative model for this type of data, which we hope will frame future analysis of CRISPR-Cas9 lineage tracing systems, akin to the Jukes-Cantor model in other molecular phylogenetic studies. With this theoretical framework and the accompanying algorithms, our work naturally complements recent efforts to develop and understand algorithms for this lineage tracing data [35]. Ultimately, we believe that this work will continue to inform and orient both algorithmic and experimental methods as the technology and field evolve.

## Acknowledgements

We acknowledge Matt Jones for valuable feedback on the introduction and discussion sections. We additionally acknowledge Satish Rao, Sebastian Prillo, and members of the Yosef Lab for helpful discussions.

## Author Contributions

RW conceived of the problem statement and led the development of the theoretical analyses. AK and RZ aided in the development of the theoretical analyses. The simulations were conceived by NY, RW, and RZ, and implemented by RZ. NY supervised the completion of the work. All authors contributed towards writing the manuscript.

## Additional Proofs

### Analysis of the *δ* Function

Since *δ*(*d*) is the only part of the bound that varies according to depth, any universal threshold to the oracle decision must take it into account. It can be verified that 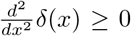 which means *δ* is convex. Setting *δ*′(*x*) = 0, we have that the minimum of *δ* occurs at:

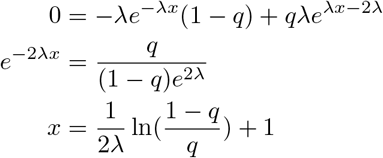

Let *x*\* be the minimum of *δ*. If *q <* 1/2, then 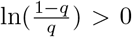, which means that *x*\* *>* 1, so on the interval [0, 1], the minimum occurs at *x* = 1, at which point *δ*(*x*) = *e*^−*λ*^. On the other hand, if *q* ≥ 1/2, then the minimum will occur at 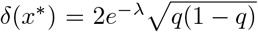 if *x*\* ∈ [0; 1] and δ(0) = 1 – *q* + *qe*^−2λ^ if *x*\* < 0. Note, *x*\* gets smaller as *q* → 1, so if *q* = 1, then the minimum is precisely *e*^−2*λ*^. Let *d*\* ∈ [0, 1] be an arbitrary depth, and let *δ*\*(*d*\*, *q, λ*) = min_*x*∈[0*,d*\*]_ *δ*(*x*). We then have:

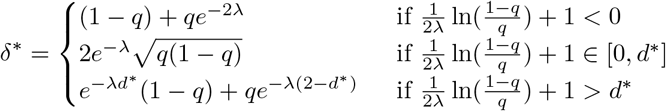

#### Proof of lemma 1

Let (*a, b*|*c*) be a triplet with *depth*(*LCA*(*a, b, c*)) = *d*, and let *α* = *dist*(*LCA*(*a, b, c*), *LCA*(*a, b*)). Let *Y* = *s*(*a, b*) and *X* = *s*(*b, c*). For a particular character *χ_i_*, let 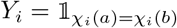 and let 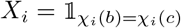. We use the following results:

##### lemma 6

*If for every character χ_i_, P* [*Y_i_* − *X_i_* = −1] *is a decreasing function of α and P* [*Y_i_* − *X_i_* = 1] *is an increasing function of α for all α* ∈ [0, 1], *then for any t, P* [*Y* − *X* ≥ *t*] *is an increasing function of α.*

**Proof:** For a given character 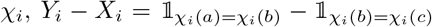 has 3 possible outcomes: {1, 0, −1}. Thus, if *P* [*Y_i_* − *X_i_* = −1] is a decreasing function of *α* and *P* [*Y_i_* − *X_i_* = 1] is an increasing function of *α* for all *α* ∈ [0, 1], then *P* [*Y_i_* − *X_i_* ≥ *t*] for any *t* is an increasing function of *α*. Stating that *P* [*Y_i_* − *X_i_* ≥ *t*] is an increasing function of *α* is identical to stating that for any *α*_1_ and *α*_2_ such that 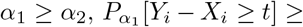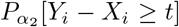. We use the known result from probability theory that for random variables 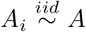 and 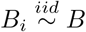 such *P* [*A* ≥ *t*] ≥ *P* [*B* ≥ *t*] for all *t*, then *P* [∑_*i*_*A_i_* ≥ *t*] ≥ *P* [∑_*i*_*B_i_* ≥ *t*]. Thus, as *Y* − *X* = ∑_*i*_*Y_i_* − *X*_*i*_ and *Y_i_* − *X_i_* is independent and identically distributed as we assume each character operates independently and identically, then 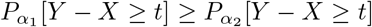. Thus, *P* [*Y – X ≥ t*] is an increasing function of *α* for any *t*.

##### lemma 7

*For every character χ_i_, P* [*Y_i_ – X_i_* = 1] *is a decreasing function of α and P* [*Y_i_ – X_i_* = 1] *is an increasing function of α for all α* ∈ [0, 1]*. Additionally, this result holds in both the general case and stochastic-only missing data cases.*

##### Proof

###### Case with no missing data

First, we examine *P* [*Y_i_ – X_i_* = 1]. *Y_i_ – X_i_* = −1 for a character *χ_i_* corresponds to that character acquiring the same mutation on both the path from *LCA*(*a, b*) to *b* and the path from *LCA*(*a, b, c*) to *c*, and not acquiring that mutation on the path from *LCA*(*a, b*) to *a*. Additionally, no mutation must be acquired at *χ_i_* on the path from the *r* to *LCA*(*a, b, c*) nor the path from *LCA*(*a, b, c*) to *LCA*(*a, b*). Thus we have:

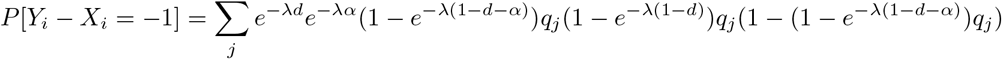

Taking only the terms that depend on *α*, we have:

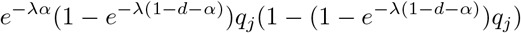

To show that this value decreases with *α*, we show that the first derivative is negative with respect to *α*. We use the following form of the derivative:

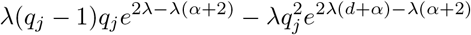

This value is positive owing to the fact that *q_j_* ∈ [0, 1] for all *j*. Thus, the function within the summation is decreasing in terms of *α*. Using the fact that the summation of decreasing functions is decreasing, the overall function is thus decreasing in terms of *α*.

Secondly, we examine *P* [*Y_i_ – X_i_* = 1]. *Y_i_ – X_i_* = −1 for *χ_i_* corresponds to a mutation occurring in both *a* and *b*, but not in *c*. A mutation can occur in *a* and *b* if it appears on the path from *LCA*(*a, b, c*) to *LCA*(*b, c*), or if it appears independently in the paths from *LCA*(*a, b*) to both *a* and *b*. Additionally, this mutation cannot appear on the path from *LCA*(*a, b, c*) to *c*, and no mutations can occur on the path from *r* to *LCA*(*a, b, c*). Thus, we have:

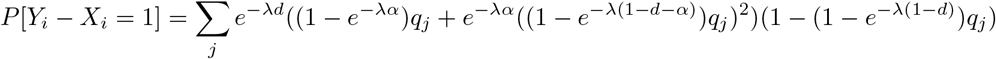

Taking on the terms that depend on *α*, we have:

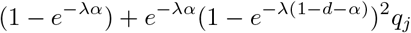

To show that this value is increasing with *α*, we show that the first derivative is positive with respect to *α*. We use the following form of the derivative:

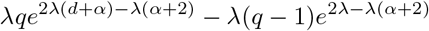

This value is positive owing to the fact that *q_j_* ∈ [0, 1]. Thus, the function within the summation is decreasing in terms of *α*. Using the fact that the summation of decreasing functions is decreasing, the overall function is thus decreasing in terms of *α*.

###### General Missing Data Case

Next, we examine the general case with both stochastic and heritable missing data. In this case, we define *s*(*a, b*) (analogously *s*(*b, c*)) as the number of characters shared by *a, b* that do not have dropout in either *a, b* or *c*. As we now simply condition on the fact that *a, b, c* must all be present, we add an additional (1 − *p_d_*) term to both *P* [*Y_i_ – X_i_* = −1] and *P* [*Y_i_ – X_i_* = −1]. As this term does not depend on *α*, both functions depend on *α* as they do in the case without missing data.

###### Stochastic-only Missing Data Case

Finally, we examine the case with only stochastic missing data. Here we define *s*(*a, b*) as the number of mutations shared by *a* and *b* in characters that did not suffer dropout in either sample. Thus, in analyzing *Y_i_ – X_i_* we must consider additional cases in which dropout in one cell can hide the fact that two cells inherited the same mutation.

First, we examine *P* [*Y_i_ – X_i_* = −1]. *Y_i_ – X_i_* = −1 for a character *χ_i_* corresponds to that character acquiring the same mutation in *b* and *c*, not acquiring dropout in neither *b* nor *c*, and either observing dropout or not observing that mutation in *a*. For this to occur, a mutation can occur on the path from *r* to *LCA*(*a, b, c*) while *a* acquires dropout, or the same mutation can occur on the path from *LCA*(*a, b, c*) to *LCA*(*a, b*) and the path from *LCA*(*a, b, c*) to *c* while *a* acquires dropout, or the mutation can occur on the path from *LCA*(*a, b*) to *b* and the path from *LCA*(*a, b, c*) to *c* while not appearing in *a*. The probability of this is:

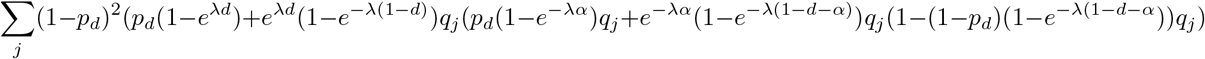

Taking the terms that depend on *α*:

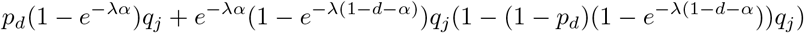

To show that this value decreases with *α*, we show that the first derivative is negative with respect to *α*. We use the following form of the derivative:

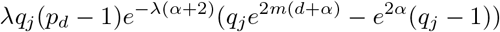

This value is positive owing to the fact that *q_j_* ∈ [0, 1] for all *j* and that *p_d_* ∈ [0, 1). Thus, the function within the summation is decreasing in terms of *α*. Using the fact that the summation of decreasing functions is decreasing, the overall function is thus decreasing in terms of *α*.

Secondly, we examine *P* [*Y_i_ – X_i_* = −1]. *Y_i_ – X_i_* = −1 for *χ_i_* corresponds to a mutation occurring in both *a* and *b*, not acquiring dropout in neither *a* nor *b*, and either observing dropout or not observing that mutation in *c*. For this to occur, a mutation can occur on the path from *r* to *LCA*(*a, b, c*) while *c* acquires dropout, on the path from *LCA*(*a, b, c*) to *LCA*(*a, b*) while the mutation is not acquired in *c* or *c* acquires dropout, or the mutation occurs independently on the path from *LCA*(*a, b*) to *a* and *b* while not appearing in *c*. The probability of this is:

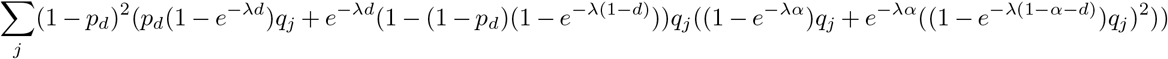

Taking on the terms that depend on *α*, we have:

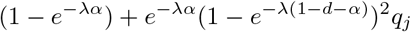

Note that this is the same value as above in the case without missing data, and hence the function will have the same dependence on *α* as in that case. Hence, the function is overall decreasing with *α*.

**Proof:** By lemmas 6 and 7, *P* [*s*(*a, b*) − *s*(*a, c*) ≥ *t*] is an increasing function of *α*. Thus, the minimum value of *α* results in the minimum value of *P* [*s*(*a, b*) − *s*(*a, c*) ≥ *t*]. This value occurs at *α* = *ℓ*\*, showing lemma 1.

#### Proof of lemma 3

Given a triplet (*a, b|c*), with LCA at depth at most *d*\*, we defined *ϵ* to be the probability that no dropout occurs in a particular character of all three cells. First we will justify the assumption that *ϵ* (1 − *p_d_*)^3^, which is to say the dropout events are positively correlated. Let *p_h_* be the probability that heritable dropout occurs on the path from *r* to *a*. Let *p_b_* be the probability that a heritable dropout occurs on the path from *LCA*(*a, b*) to *b* given that no dropout has occurred on the path from *r* to *LCA*(*a, b*) and define *p_c_* similarly. Note that *p_b_ ≤ p_c_ ≤ p_h_* since the probability a dropout occurs along a path increases with the length of the path. Let *p_s_* be the probability that a stochastic dropout occurs at given character on a leaf given that no heritable dropout has occurred yet on that character. Then we have *ϵ* = (1 − *p_h_*)(1 − *p_b_*)(1 − *p_a_*)(1 − *p_s_*)^3^ ≥ ((1 − *p_h_*)(1 − *p_s_*))^3^ = (1 − *p_d_*)^3^. Since at least one of the cells in the triplet needs to not incur dropout at a character in order for all three of them to have no dropout, we also have *ϵ* ≤ 1 − *p_d_.*

To prove the bounds on *k* we will proceed as in the proof of lemma 2 and assume that *dist*(*LCA*(*a, b, c*), *LCA*(*a, b*)) = *ℓ*\*, noting that lemma 7 extends the result of lemma 1 to the general case with missing data. Let *Y* = *s_c_*(*a, b*) and *X* = *s_a_*(*b, c*). Thus, we have:

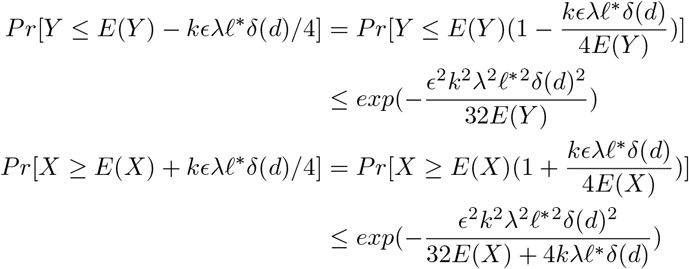

Since *E*(*Y*) ≤ *kϵe*^−*λd*^*λℓ*\* and *E*(*X*) ≤ *kϵe*^−*λd*^*λ*^2^*q*, we see that both probabilities are at most *n*^−3^*ζ* when

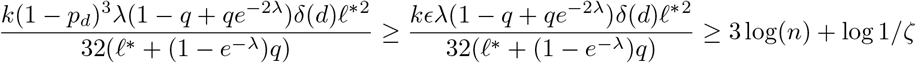

If both bad events don’t happen, then we have *Y – X ≥ kϵλℓ*\**δ*\*/2 *k*(1 − *p_d_*)^3^*λℓ*\**δ*\*/2 = *t*. This gives the necessary bound for condition *i*) to hold.

To get guarantees on the second condition, let *X* = *s_b_*(*a, c*) for any triplet *a, b, c* whose LCA has depth at most *d*\* and where *c* is the outgroup. we have that

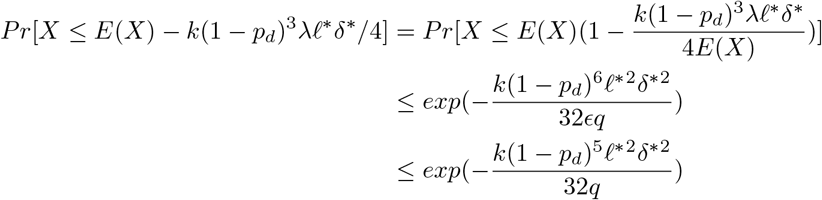

To ensure that this probability is at most *n*^−3^*ζ*, it suffices to take

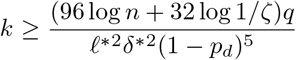

The rest of the argument is exactly the same as in the proof of lemma 2.

#### Proof of lemma 4

For a triplet (*a, b*|*c*) in the case without dropout, any mutation that occurred on the path from the root to *LCA*(*a, b, c*) is inherited by Each member of the triplet, and thus *s*(*a, b*) – *s*(*b, c*) = *s_c_*(*a, b*) – *s_a_*(*b, c*). But in the case of missing data, this is no longer true as mutations that occurred before *LCA*(*a, b, c*) may be obscured by dropout and therefore not present in the character information of *a*, *b*, or *c*. We must now account for these early mutations in our calculations.

Let *p_s_* be the stochastic missing data rate and let *s*(*a, b*) be the number of mutations shared between *a* and *b*, ignoring characters that have dropout in either *a* or *b*. The number of mutations shared by *a, b* after their divergence is now Binomial(*k, p*) where *p* is

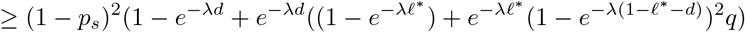

Here (1 − *p_s_*)^2^ is the probability that this character does not acquire dropout in neither *a* nor *b*, 1 − *e*^−*λd*^ is the probability that a given mutation occurred before *d*, and of the remaining terms the left term is the probability the mutation occurred on the path from *LCA*(*a, b, c*) to *LCA*(*a, b*), and the right term is the probability a given mutation is shared by *a, b* due to convergent evolution.

Thus:

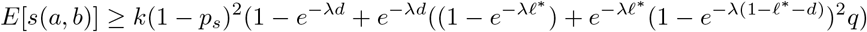

Similarly:

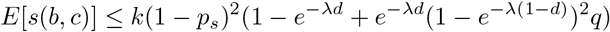

Thus, we have that:

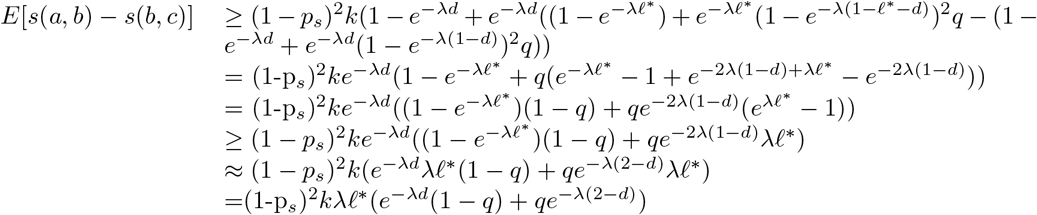

Again taking *δ*(*d*) = *e*^−*λd*^(1−*q*)+*qe*^−*λ*(2−*d*)^. We then have that for any triplet (*a, b*|*c*), where *depth*(*LCA*(*a, b, c*)) = *d* and *dist*(*LCA*(*a, b, c*), *LCA*(*a, b*)) ≥ *ℓ*\*

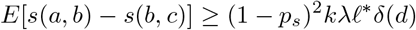

First, we will show that condition *i*) will hold with probability 1 − *ζ* if:

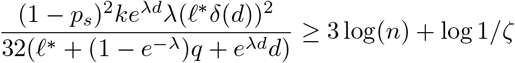

To see this, let (*a, b|c*) be any triplet at depth at most *d*\* and the distance between their LCAs be at least *ℓ*\*. By lemma 1, we can WLOG assume that *dist*(*LCA*(*a, b, c*), *LCA*(*b, c*)) = *ℓ*\* because that is the worst case, i.e. the case where *P* [*s*(*a, b*) – *s*(*b, c*) ≥ *t*] is minimized. Note that lemma 7 extends the result of lemma 1 to the case with only stochastic missing data. Any condition sufficient for this case will be sufficient overall. Let *Y* = *s*(*a, b*) and *X* = *s*(*b, c*). Since *E*(*Y*) – *E*(*X*) ≥ (1 − *p_s_*)^2^*kλℓ*\**δ*\* = 2*t*, in order to ensure that *Y − X ≥ t*, it suffices to have *Y > E*(*Y*) = *t/*2 and *X < E*(*X*) = *t/*2. To show that both occur with high probability, we have:

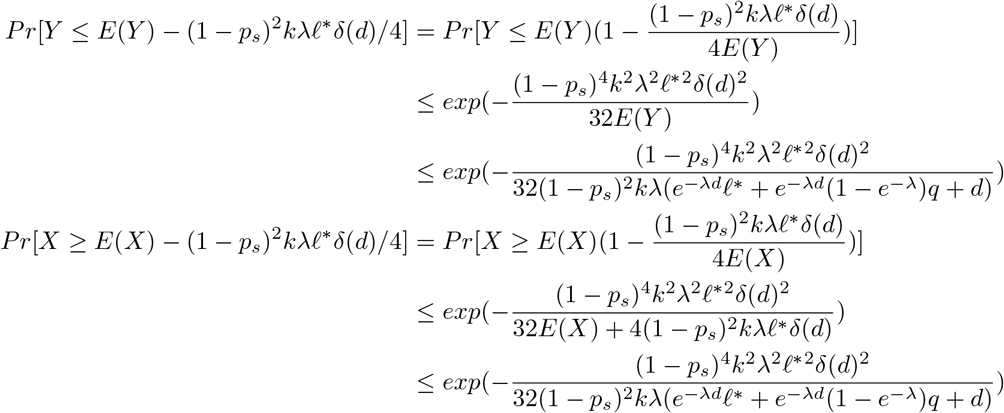

The last line follows from the fact that *δ*(*d*) ≤ *e*^−*λd*^ and *E*(*X*) ≤ (1 − *p_s_*)^2^*ke*^−*λd*^(1 − *e*^−*λ*^)^2^*q* + *λd* ≤ (1 − *p_s_*)^2^*ke*^−*λd*^(1 − *e*^−*λ*^)*λq* + *λd*. Since *e^λd^δ*(*d*) = 1 − *q* + *qe*^−2*λ*(1−*d*)^ ≥ 1 − *q* + *qe*^−2*λ*^ and any *d < d*\*, in order to ensure that both bad events have probability at most *ζn*^−3^, it suffices to take

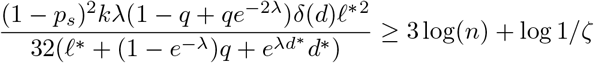

Applying the same argument to *s*(*a, b*)−*s*(*a, c*) and combining both results gives *P* [*s*(*a, b*)−max(*s*(*a, c*), *s*(*b, c*)) *< t*] ≤ 4ζ*n*^−3^.Taking a union bound over all 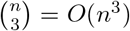 triplets, we see that the probability of condition *i*) failing for any triplet is at most ζ.

To get guarantees on the second condition, note that condition *i*) implies that condition *ii*) holds for all triplets separated by an edge of length at least *ℓ*\*. Thus we can focus on the triplets that are not covered by condition *i*). Let (*a, b|c*) be an arbitrary triplet such that *dist*(*LCA*(*a, b, c*), *LCA*(*a, b*)) *< ℓ*\* and again let *Y* = *s*(*a, b*), *X* = *s*(*b, c*) and *d* be the depth of *LCA*(*a, b, c*) (note that we are focusing WLOG on *s*(*b, c*) since it has the same distribution as *s*(*a, c*)). We want to show that with high probability, *X – Y < t*. Again, it suffices to upper bound *P* [*Y ≤ E*(*Y*) − *t/*2] and *P* [*X ≥ E*(*X*) ≥ *t/*2] because *E*(*Y*) ≥ *E*(*X*). Note that we have already bounded the second quantity. To bound the first quantity, note that the worst case scenario is that *dist*(*LCA*(*a, b, c*), *LCA*(*a, b*)) is as small as possible, but since this quantity can be arbitrarily small, we can assume that in the worst case, *dist*(*LCA*(*a, b, c*), *LCA*(*a, b*)) = 0, which means *Y* has the same distribution as *X*. Note that this case technically cannot happen as it would imply that 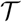 has a trifurcating branch but it is possible to get arbitrarily close to this case with no restrictions on edge lengths. This gives:

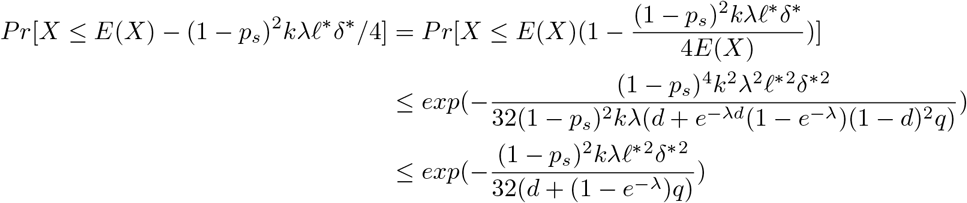

Thus, if we take:

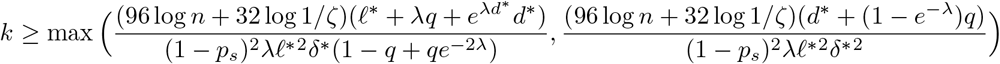

then we have *P*[*X* –*Y* ≥ *t*] < ζ*n*^−3^. By symmetry, this means *P*[*s*(*b*, *c*) – *s*(*a*, *b*) ≥ *t* ⋃ *s*(*a*, *c*) – *s*(*a*, *b*) ≥ *t*] ≤ 2*ζn*^−3^ Since we can union bound over one bad event of probability at most 2*ζn*^−3^ for each of the 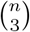 triplets, we have that conditions *i*) and *ii*) both hold with probability at least 1 − *ζ*.

#### Proof of lemma 5

For this proof, we use the following results:

##### lemma 8

*Let ρ be the maximum edge length in* 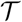 *. For any character, state pair* (*i, j*), *if there exists a number p >* 0 *which satisfies* ((1 − *e*^−*λρ*^)*q_j_* + *p*)^2^ ≤ *p, then p is an upper bound on P_i,j_* (*v*) *for any node 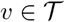.*

**Proof:** We will proceed by induction on 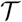. Suppose *v* is a leaf. Then *P_i,j_* (*v*) = 0, since if the *i^th^* character does not mutate, it cannot take on state *j*. Now let *v* be an arbitrary non-leaf vertex, with children *u* and *w*. By our inductive hypothesis, *P_i,j_* (*w*) ≤ *p* and *P_i,j_* (*u*) ≤ *p*. Since the length of the edge from *v* to either of it’s children is at most *ρ*, the probability that the character mutates to state *j* on either edge is at most 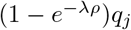. Thus, if we condition on the fact that *χ_i_* is not mutated on *v*, we have:

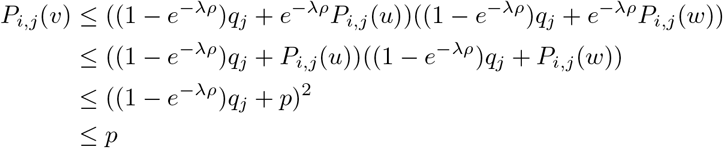

Where the last inequality follows from our assumption on *p*. Given the above lemma, we can simply solve for *p* to find an upper bound on all *P_i,j_*(*v*) in 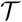.

##### lemma 9

*Suppose the maximum edge length satisfies:*

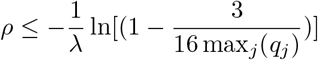

*Then we have* 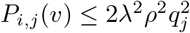 *for any node* 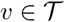

**Proof:** Let *y* = (1 − *e*^−*λρ*^)*q_j_*. Note that by our above assumption, *y* ≤ 3/16. By our above assumption, we have:

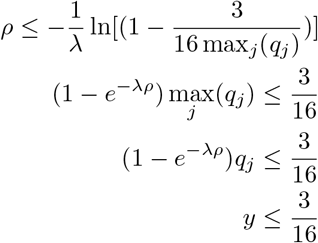

Next, by lemma 8, we know that if there is a *p >* 0 such that (*y* + *p*)^2^ ≤ *p*, then such a *p* would be an upper bound on all *P_i,j_*(*v*). We can find such a *p* by setting the inequality to an equality and finding the smallest root of the resulting polynomial.

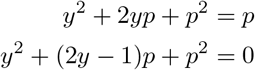

Our initial assumption of *ρ* guarantees that 4*y <* 1, which means the smallest root of the polynomial above can be given as

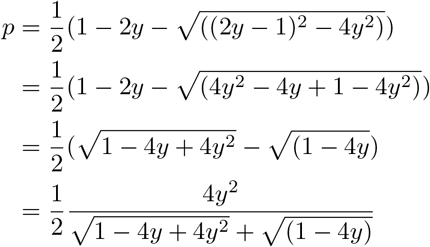

Where the last line follows by multiplying the numerator and denominator by 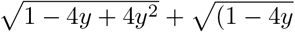. To upper bound *p*, we have

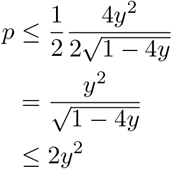

Where the last inequality follows from the fact *y* ≤ 3/16, which means the denominator is at least 1/2. Finally, we have 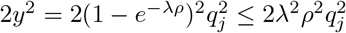.

Now simply take 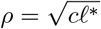 and apply lemma 9 to show lemma 5.

### Alternative Analysis of the Bottom-up Algorithm

#### Theorem 4

*The constraint on λ and q in Theorem 3 can be replaced with*

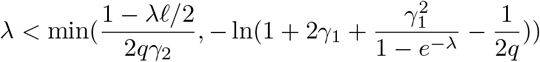

*Where* 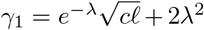 *and* 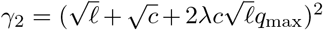. *In that case, the Bottom-Up Algorithm returns the correct tree with probability at least* 1 − *ζ if*

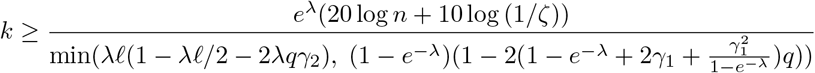

Note this means that as *ℓ* → 0, our constraint on *λ* and *q* becomes 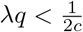 and our bound on *k* approaches

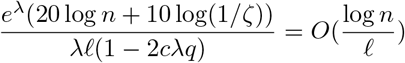

**Proof of Theorem 4:**. To upper bound the probabilities of the bad events, it suffices to ensure that *E*(*Z*) *> E*(*X*) and bound the quantity 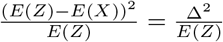. Since this quantity depends on *α*, we will first derive a lower bound on this quantity and determine where the minimum occurs for *α* ∈ [*ℓ,* 1] as follows:

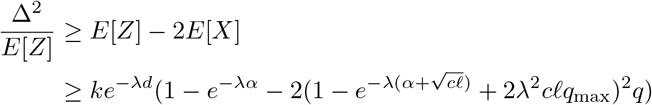

Now let 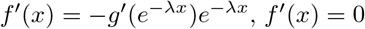. Next we will show that for any interval [*a, b*], the minimum of *f* on [*a, b*] occurs at either *a* or *b*. Let *y*(*α*) = *e*^−*λα*^, and let 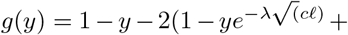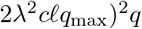. Then *f* = *g* ◦ *y*. Note that *g*′ is linear with negative slope, as

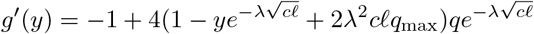

Since *f* ′(*x*) = −*g*′(*e*^−*λx*^)*e*^−*λx*^, *f* ′(*x*) = 0 only when *g*′(*e*^−*λx*^) = 0. Since *e*^−*λx*^ is an increasing function, there can be at most one point where *f* ′ is 0. On the other hand, one can verify that lim_*x*→−∞_ *f* ′(*x*) = ∞, which means that if there is any *x* where *f* ′(*x*) = 0, then *f* ′ is positive on (∞, *x*) and negative on (*x,*), which means *x* must be a local maximum. Thus, any minimum of *f* on an interval [*a, b*] can only occur on the boundaries.

*α* = 1 **case:**

In the case where *α* = 1, our lower bound can be written as

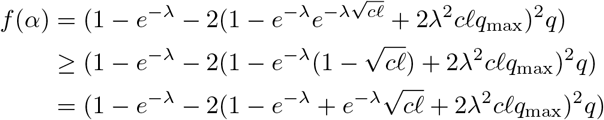

To make the notation easier to follow, let 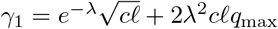. Then we have

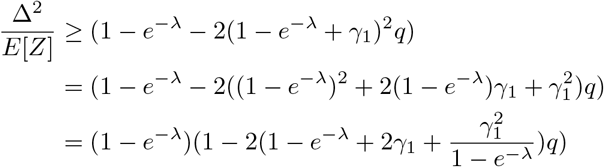

Thus, in order for the bound to be non-trivial, we need 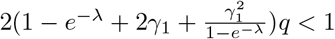, which means

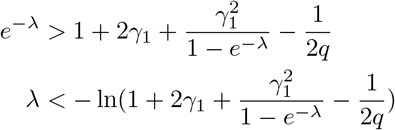

Note that as *ℓ* → 0, the RHS approaches 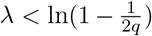 which is at least 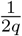, and in general, when *λ* satisfies the constraint above, *f* (1) is lower bounded by a constant independent of *ℓ*. Also, note that the bound is bound is trival if the term inside the ln is negative.

*α* = *ℓ* **case:**

In this case, our lower bound becomes as follows:

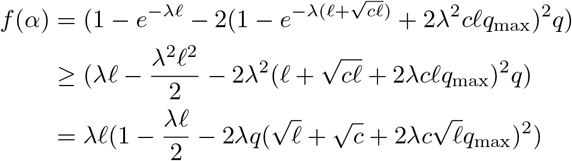

Now let 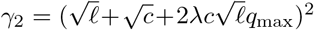. Then in order for the bound to be non-trivial, we need 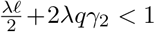, which means

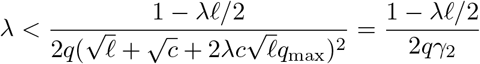

Note that as *ℓ* → 0, the RHS becomes 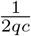. Since 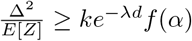, when the constraints:

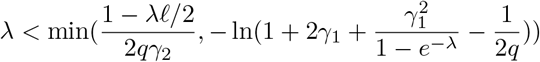

are satisfied, and if we take

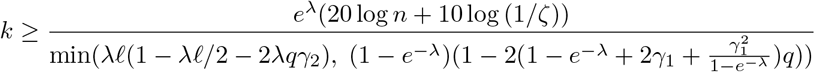

the probability that either of the above events occur is at most *n*^−2^*ζ*. In other words, for any pair of vertices *u, w* that are not children of the same node, there will be a vertex *u*′ that such that *LCA*(*u, u*′) is a descendent of *LCA*(*u, w*) and *P* [*s*(*u, u*′) ≤ *s*(*u, w*)] ≤ 2*n*^−2^*ζ*. If *s*(*u, u*′) *> s*(*u, w*), then (*u, w*) cannot be the first pair of incorrectly joined vertices. Taking a union bound over at most *n*^2^/2 pairs of vertices, we see that with with probability at least 1 − *ζ*, there is no first pair of incorrectly joined vertices, which means the algorithm is correct.

Since the bound on *f* (*ℓ*) depends on *ℓ*, it is general much smaller than the bound on *f* (1), which is lower bounded by a constant. Thus, asymptotically, the bound on *k* is

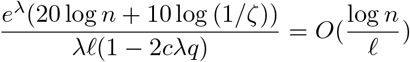

## Description of Simulations and Scoring Criteria

Simulations and algorithms are implemented in Python in Cassiopeia software suite [10] (https://github.com/YosefLab/Cassiopeia), utilizing the NetworkX package [45].

### Implementation of Algorithms

#### Threshold Algorithm

Due to ties in the number of shared characters, sometimes the edge-removal procedure produces more than two connected components on the sample graph *G*. If this occurs, we enforce a bifurcation in the tree by merging the components *C*_1_, *C*_2_, *…, C_n_* into two groups. We expand the previous pseudocode as follows:

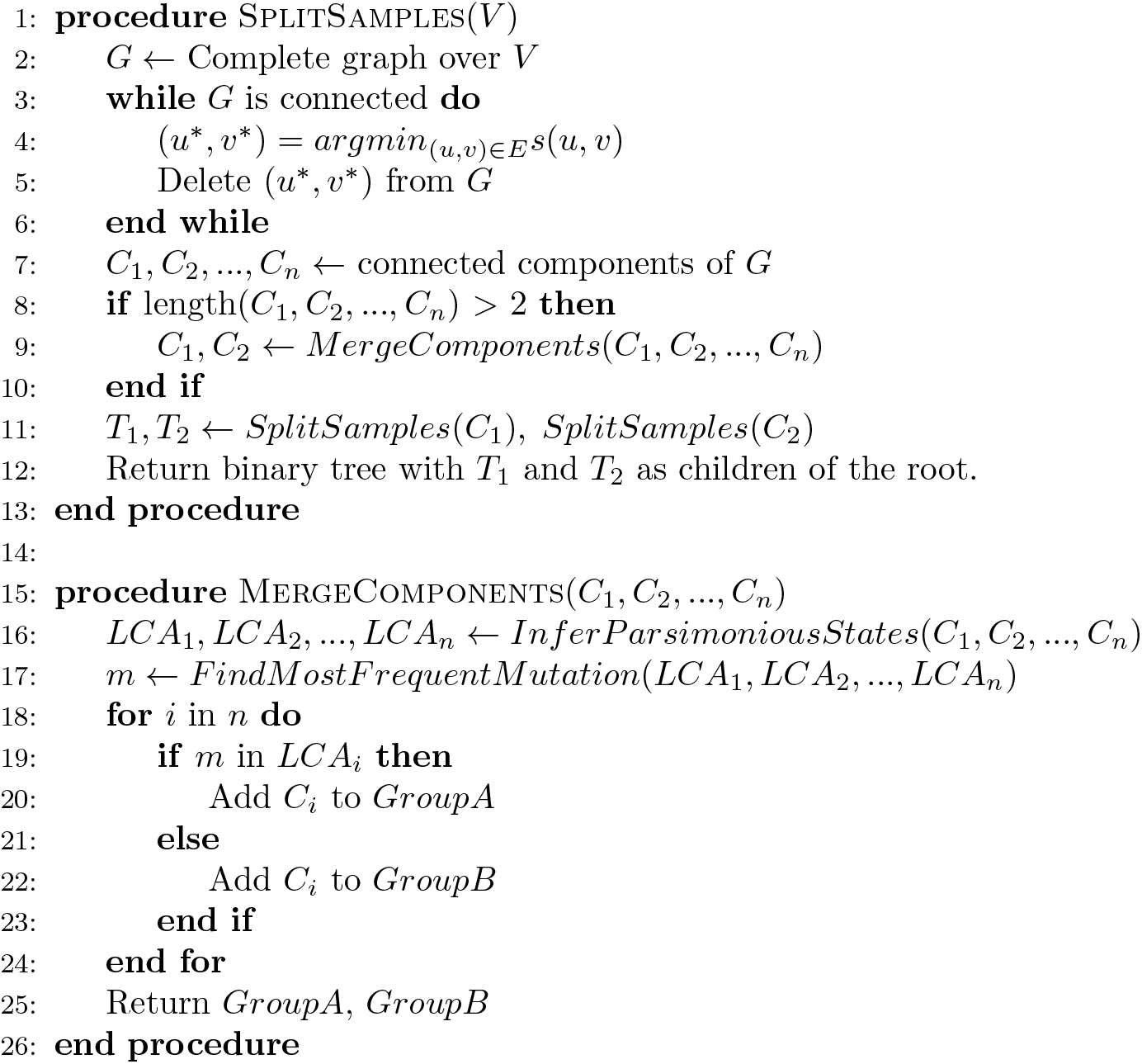

Here, we use a naive parsimony approach. We infer the character states of the LCA of each component as the most parsimonious states given the states of the leaves in that component (a mutation appears in the LCA of a component only if it shared by all samples in that component, discounting samples with missing data at that character). Then, the mutation shared by the most LCAs is found. Heuristically, this mutation occurred early in the phylogeny and thus components sharing this mutation are more closely related. Thus, we separate components into two groups based on whether or not the LCA has this mutation.

This algorithm is implemented as the “PercolationSolver” class in the “solver” module of the Cassiopeia codebase. Here, the default arguments are used, with “joining solver” specified as an instance of the “VanillaGreedySolver” class.

#### Bottom-Up Algorithm

The implementation of the Bottom-Up Algorithm follows the description in the main text. If a tie occurs in the number of shared mutations between nodes, then an arbitrary pair is chosen to be merged first.

This algorithm is implemented as the “SharedMutationSolver” class in the “solver” module of the Cassiopeia codebase. Here, the default arguments are used.

### Simulating Lineage Tracing Experiments

In our simulations, we simulated forward-time lineage tracing experiments using our generative model. We split the simulation into two steps.

#### Simulating Cell Division Topologies

First, we simulate a continuous-time, binary, symmetric cell division topology. Then, we simulate CRISPR-Cas9 lineage tracing data over the given topology. The end result is a phylogenetic tree representing the single-cell lineage tracing experiment. The tree topology also records the ground truth phylogenetic relationships between the observed cells.

We begin by describing the two simulation schemes used for the tree topology. The first scheme simulates a cell division regime with regular cell division (uniform edge lengths). We start with a complete binary tree and add an implicit root, attaching this root to the root of the complete binary tree by an edge. The edge represents the lifetime of the root along which mutations can be acquired. For all figures besides Figure 6, we generated trees with 256 leaves representing cells observed at the end of the experiment. For Figure 6, we generated trees of various sizes, exponentially increasing *n*. Given that the number of edges in the path from the implicit root to each leaf has *log*(*n*) + 1 edges, each edge has uniform length, and the length of the experiment is normalized to one, each edge has length 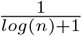.

The second simulation scheme represents an asynchronous cell division regime, with stochastic waiting times between cell divisions and cell death. We model a forward-time Bellman-Harris model with extinction [46]. This generalizes the birth-death process [47], a commonly used phylogenetic model, such that the distribution of waiting times between division and death events are arbitrary. In our case, waiting times between division events are distributed according to an exponential distribution that is shifted by a constant *a* = 0.05, representing minimum time between cell division events. The distribution of death waiting times is distributed exponentially, as we assume that cell death does not have a minimum time. The stopping condition is when all lineages reach time = 1, meaning that each lineage will have total path length from the root of 1. We present the pseudocode used for this simulation here:

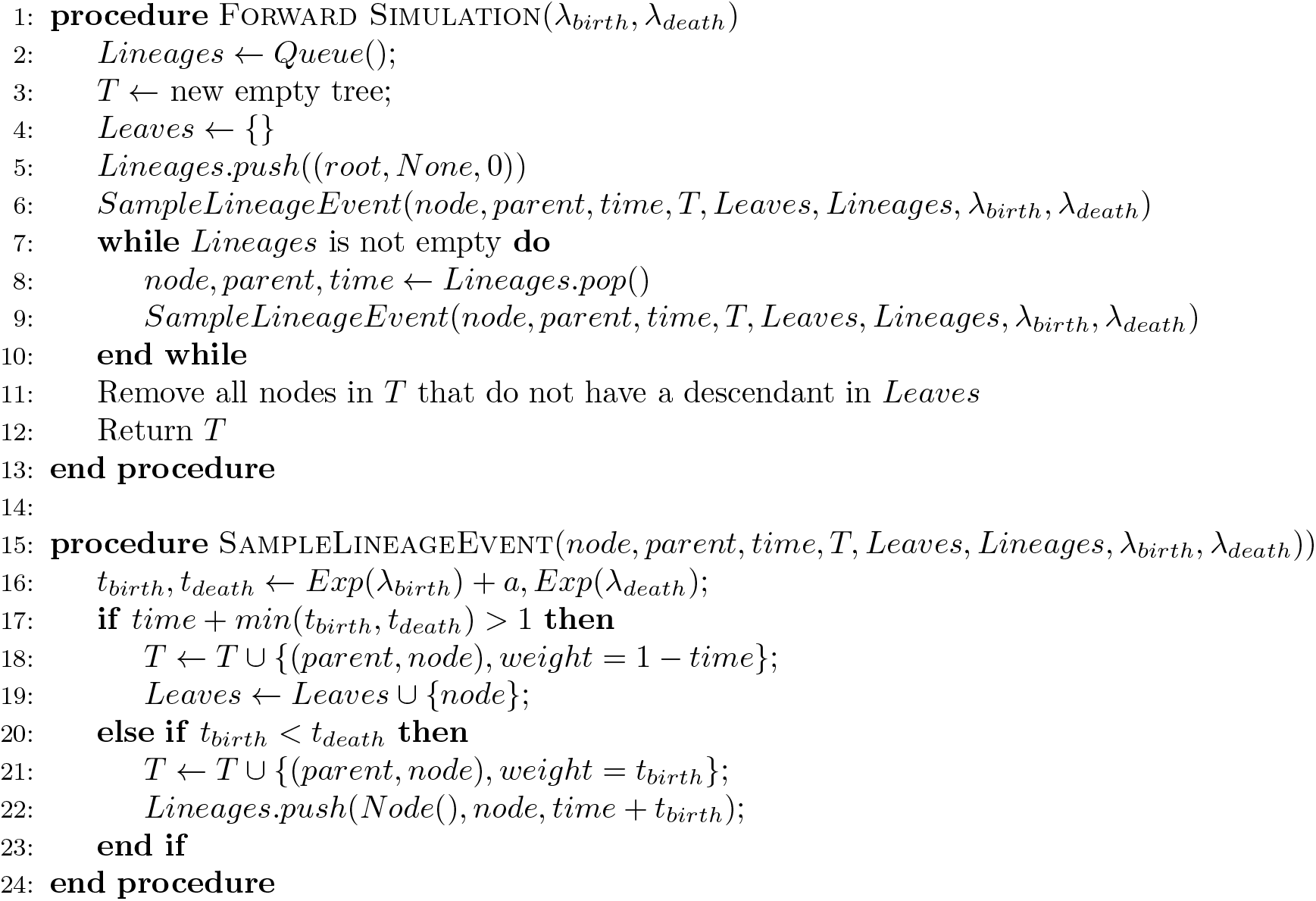

To control for the number of leaves in the simulated trees, we take only trees that have between 205 and 307 leaves (if the procedure terminates with no living lineages, then we consider that tree to have 0 leaves). Note that in the trees generated by this process sister nodes need not have the same edge length and the root will have a singleton edge as in the binary case along which mutations can occur. We chose rates for the division and death waiting distributions (23.70 and 2.12, respectively) that gave an average of around 256 leaves over 1000 simulations. These rates were chosen assuming that the rate of division was 10 times that of the rate of death, and then correcting the death rate to increase the mean waiting time by *a* = 0.05 to match the shift in mean in the distribution of division times.

Due to the stochastic nature of this division process, we cannot exactly control for *ℓ* if we stop the experiment at a specified time. This is due to the fact that edges at the leaves of the tree may be very small if the stopping criterion is reached before the length can reach *a*, thus making the minimum edge length in the tree technically potentially smaller than *a*. We contend that these edges should not impact the analysis though. Small edges make it difficult to discern which neighboring clades are actually closer in relation. These small edges only occur at the bottom of the tree though and would only affect the edge lengths leading to single leaves, which are trivially discerned as a cherries with their neighbors, meaning that *ℓ* would still effectively be 0.05 in this case.

This topology simulation framework is implemented in the Cassiopeia codebase as the “BirthDeathFitness-Simulator” class in the “simulator” module. Here, “birth waiting distribution” is set to a lambda function that takes a rate and returns a random waiting time from an exponential distribution with that rate, shifted by 0.05. “initial birth scale” is set to ≈ 23.70. “death waiting distribution” is set to a lambda function that takes no arguments and returns a random waiting time from an exponential distribution with that rate 1/ ≈ 23.70 + 0.05. “experiment time” is set to 1. The other arguments are set to their defaults.

#### Simulating CRISPR-Cas9 Lineage Tracing Data

Given a tree topology, we simulate a CRISPR-Cas9 mutagenesis process over it. Along each edge with length *t*, independently for each character, we simulate the probability that a mutation will occur as 1 − *e^λt^*. If a character has been chosen to mutate, we then draw from the state distribution to determine the state the character acquires. In this case, this is a uniform distribution with *m* = 1*/q* states (note that in the uniform case, 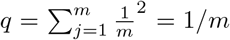. Once a mutation is acquired on an edge, that mutation persists in all downstream nodes. Then the mutations acquired along the path from the implicit root is maintained for each leaf node, which then forms the observed character information for all observed cells. If the simulation involves missing data, then *p_s_* proportion of characters are uniformly at randomly changed to a state representing missing data (−1). This character information is the input to the reconstruction algorithm.

This lineage tracing simulation framework is implemented in the Cassiopeia codebase as the “Cas9LineageTracingDataSimulator” class in the “simulator” module. Here, “number of cassettes” is set to the respective *k*, “size of cassette” is set to 1, “mutation rate” is set to the respective *λ*, “state priors” is set to a dictionary representing the *q* distribution, “heritable silencing rate” is set to 0, and “stochastic silencing rate” is set to 0 and 0.1, depending on whether the particular simulation has missing data. All other arguments are set to default.

### Finding the Minimum Necessary *k*

#### Exploring the Space of *k*

Here we describe the process by which we determined the minimum *k* necessary for 90% probability of a given criterion in our simulations, either full reconstruction, partial reconstruction, or triplets correct. For a given value of *k* and a given a set of parameters, we verify if it is sufficient for 90% probability of full reconstruction as follows: we simulate 10 ground truth trees, reconstruct each tree from its observed cells (leaves) using the relevant algorithm, and comparing the corresponding reconstructed tree to each ground truth tree. If ≥ 9 out of those 10 trees meet a scoring criterion, then we say that this *k* is sufficient. To alleviate the effect of noise, if 7-8 out of 10 trees meet the criterion, then we construct 20 additional trees and say *k* is sufficient if ≥ 18 of those trees meet the criterion.

To efficiently Explore the space of *k*, we first exponentially (base 2) increase the value of *k* until a max value is reached (4098 in the case of no missing data and 16384 in the case of missing data). Once we find a sufficient *k*, we perform a binary search in the bin between that value and the value before it. Finally, we record the number of trees correctly reconstructed out of 10 for each value of *k* in the binary search and perform a logistic regression on these data points. We report the value of *k* that first reaches 90% reconstruction probability predicted by the logistic regression. If no *k* is sufficient up to the max value, then we deem that the necessary value of *k* is too large for our simulations to discern and we report a missing value. To calculate the point-wise confidence intervals used in Figures 2, 4, 6 for each regression on a set of parameters, we calculate the upper and lower bounds of the 95% confidence interval from the regression coefficients using the delta method. Then, we take the upper bound on the necessary *k* as the first *k* where the lower bound exceeds 90%, and we take the lower bound as the first *k* where the upper bound exceeds 90%.

#### Full reconstruction

We say the reconstructed tree achieves perfect reconstruction if it has a Robinson-Foulds Distance of 0, meaning the trees are isomorphic with regard to their labels.

Robinson-Foulds Distance is implemented in the codebase as the “robinson foulds” method in the “critique” module. This method makes use of the the Ete3 package [48].

#### Partial Reconstruction

To determine the sufficient *k* needed for exact partial reconstruction for triplets whose LCA is up to depth *d*, we use the same framework as in the case of full reconstruction but we change the scoring criterion. We can no longer compare ground truth and reconstructed trees by Robinson-Foulds Distance, which compares the entire tree. We instead present and show the correctness of an algorithm to determine if all triplets in a tree up to depth *d* are resolved correctly in the reconstructed tree. The algorithm is as follows. Let 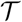 be the ground truth tree, 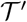 be the reconstructed tree, and *d* be the depth:

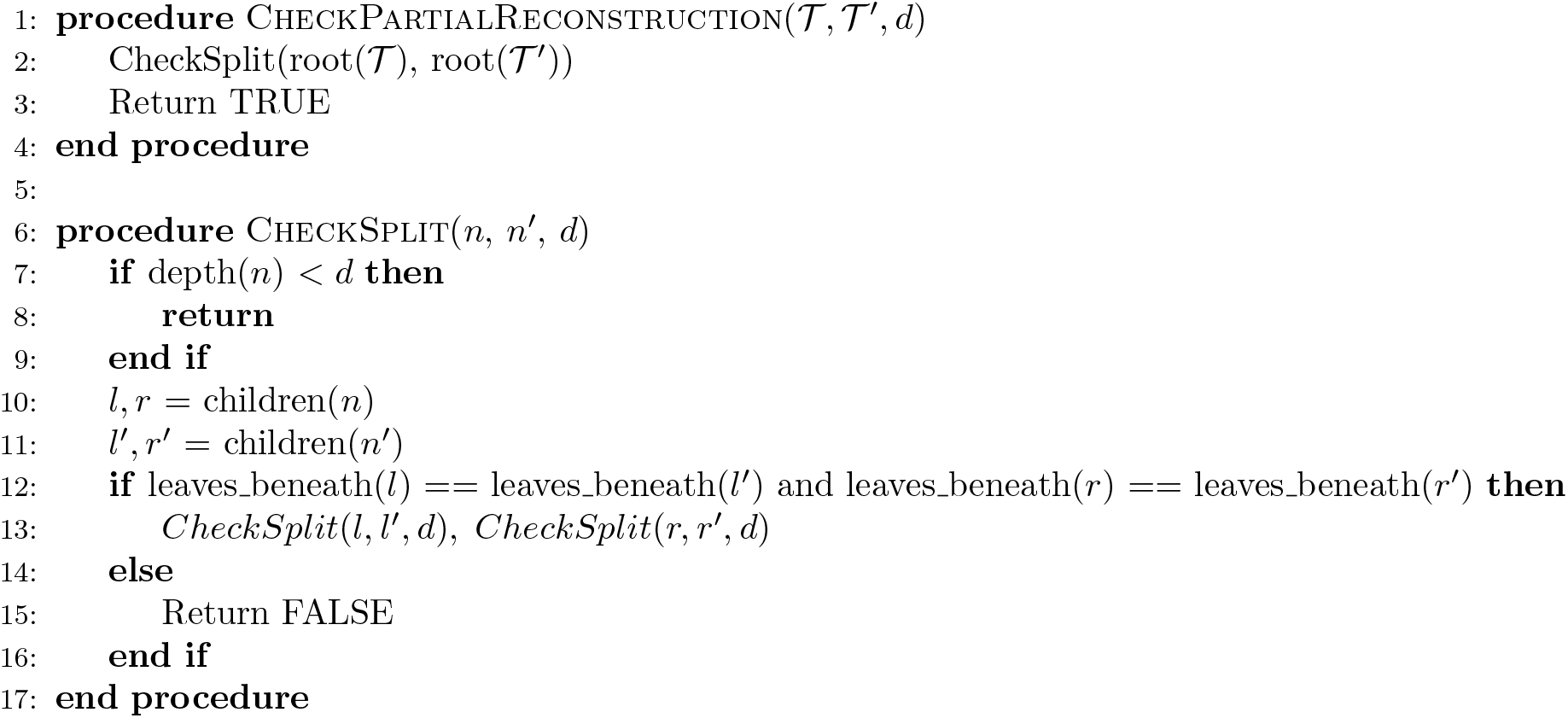

#### Claim

All triplets whose LCA is at depth *< d* 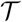 in are resolved correctly in 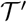 iff for every node *n < d* in 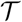 there exists a node *n*′ in 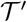 whose daughter clades partition the set of leaf descendants of that node in the same way.

**Proof: if:** For a triplet (*a, b|c*) whose LCA is node *n*, *a, b* must be in the clade of one daughter of *n* and *c* on the other. If there exists a node *n*′ in 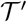 that partitions the leaf nodes into the same two clades, then for each triplet whose LCA is at *n*, *a, b* will be grouped together in the same clade with *c* on other side, hence every triplet will be resolved correctly. If there exists *n*′ for each *n < d*, then all triplets with LCA *< d* will be resolved correctly. **only if:** If there is a node *n < d* in 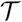 with no node *n*′ in 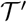 with an analogous partition, this implies that there is some partition of the leaves in 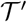 starting from the root at or above *n* that does not match the partition in 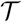, as if all partitions were correct then *n*′ must exist. If there is a non-matching partition, there is some partition {*a*_1_, *…, a_m_*}|{*b*_1_, *…, b_n_*} in 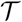 where in 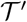 there is at least one incorrect member in one of the partitioned sets: 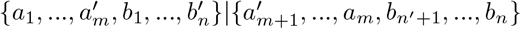. Then in 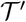, WLOG *b*_1_ is closer to some *a_i_* than some *b_i_*, and all triplets involving *b*_1_ and *a_i_* are incorrect in 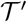. As the partition with the non-analogous partition is at depth *< d* in 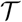, then some triplets whose LCA is at depth *< d* in are incorrectly resolved in 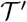.

The algorithm will find *n*′ for every *n < d* by matching it to a node in 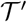 that has the same partition. If the algorithm does not find *n*′ for a certain *n < d*, then it does not exist as if all partitions checked by the algorithm match up to and including *n* in 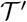 then it must exist, and the given partition cannot be formed later as leaf descendant sets cannot add members down the tree.

#### Triplets Correct

Additionally, we report the necessary *k* needed for 95% triplets correct in simulation. The triplets correct score is determined by sampling 500 triplets uniformly from the ground truth tree and counting the proportion of triplets resolved accurately in the reconstructed tree.

## Additional Simulations

### Visualization of the (*ℓ*\*, *d*\*)−Oracle

The (*ℓ*\*, *d*\*)–Oracle presented above attempts to determine the outgroup of a triplet using the difference in the number of shared mutations between its members. In Figure 5 we visualize how well the decision rule holds for correctly and incorrectly resolved triplets. We plot *s*(*a, b*) – max(*s*(*a, c*), *s*(*b, c*)) against *dist*(*LCA*(*a, b*), *LCA*(*a, b, c*)) for triplets (*a, b, c*) on simulated trees. The blue points show the difference between the mutations shared by the ingroup versus the mutations shared by the outgroup, and the orange points show the difference between the mutations shared by the outgroup and the ingroup. We see that 10% of triplets are such that *s*(*a, b*) – max(*s*(*a, c*), *s*(*b, c*)) ≤ *t* with (*a, b*) as the ingroup and thus violate condition *i*), and that 2.3% are such that *s*(*a, b*) – max(*s*(*a, c*), *s*(*b, c*)) ≤ *t* with (*a, b*) as the outgroup. Note that as *s*(*a, b*) – max(*s*(*a, c*), *s*(*b, c*)) ≤ *s*(*a, b*) − *s*(*a, c*), requiring that *s*(*a, b*) – max(*s*(*a, c*), *s*(*b, c*)) *< t* is a slightly weaker condition than condition *ii*) and thus at most 2.3% of triplets violate it. For the set of parameters used here, *t* is chosen such that the probability that a triplet is indeterminable is low and the probability that the outgroup is given incorrectly is low, showing that the oracle is relatively accurate on this regime for the low number of characters (*k* = 10). For for exact reconstruction of a tree though, we require that all triplets on that tree be separated by the threshold *t*.

As *dist*(*LCA*(*a, b*), *LCA*(*a, b, c*)) increases, this difference in the number of shared mutations between the ingroup and the non-ingroup pairs grows, making the triplets more separable. This gives us the V-shape in the figure. As the distance increases, the number of triplets that cross the threshold and thus violate either condition decreases, and after a certain point no triplets cross the threshold. This illustrates that in the result in Theorem 2 that, as we are only interested in triplets where *dist*(*LCA*(*a, b*), *LCA*(*a, b, c*)) *> ℓ*\* for larger values of *ℓ*\*, them the bound on the number of characters *k* needed to separate these triplets decreases.

### Dependence on *n*

To compare the asymptotic dependence of *k* on *n* in the theoretical bounds with the dependence of the necessary *k* in the empirical case, we simulate varying trees of size. In Figure 6 we plot the values of *k* in simulation against the bounds for both algorithms, in a regime where *ℓ* = 0.5, *q* = 0.05. From this figure it can be seen that the bounds are tight (within a constant factor) against the empirical values and share the same shape in the dependence of *k* on *n*. These results offer empirical validation of the asymptotic dependence of *k* in the bounds for both algorithms.

### Missing Data

Here we present simulations for the minimum *k* to give 0.9 probability of exact reconstruction for the case of stochastic missing data only (lemma 4). The simulations are performed with uniform edge lengths and use *p_s_* = 0.1 proportion of stochastic missing data. Visualizing the bounds for lemma 4 show that indeed higher values of *k* are necessary to overcome the lost information (Figure 7A), consistent with the additional (1 − *p_d_*)^2^ term in the denominator and gap term of *e^λd*^ d*\* when compared to the bounds of Theorem 1. Notably, the reconstruction now becomes increasingly intractable for high values of *λ*, due to the gap term being exponential in *λ*. In simulation (Figure 7B) we see that the necessary *k* is higher across the board, especially for values with high *λ*, validating the theoretical trends.

Surprisingly, unlike in lemma 2, the bound on *k* in lemma 4 has no asymptotic dependence on *q*. Taking *q* to be arbitrarily small (or even *q < ℓ/λ*) causes the bound in lemma 2 to become 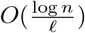, yet an arbitrarily small *q* causes the bound in lemma 4 to remain in 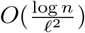. Examining the difference in the bounds between lemma 4 and lemma 2 (Figure 7C (Top)), we see that the difference grows larger in regions where *q < ℓ/λ*, indicating that the bound changes asymptotically in these regions. This same pattern is reflected in the empirical results, validating this change in dependence on *q* (Figure 7C (Bottom)).

### Triplets Correct

Previous benchmarking works do not use exact reconstruction as a metric for the accuracy of phylogenetic reconstructions. A common, more relaxed metric is the triplets correct metric (or the closely related triples distance), which measures the proportion of sampled leaf triplets that are correctly (incorrectly in the case of triplets distance) inferred by the reconstructed tree [10, 49, 35]. We present the minimum *k* necessary in simulation for high probability of a high (≥95%) triplets correct score on uniformly sampled triplets, showing the necessary *k* when exact reconstruction is not required (Figure 8). We see that the empirical necessary *k* decreases substantially overall compared to the case of exact reconstruction for both algorithms, showing that these algorithms can perform well in practice with a low number of characters according to traditional standards of accuracy.

**Figure 8:**
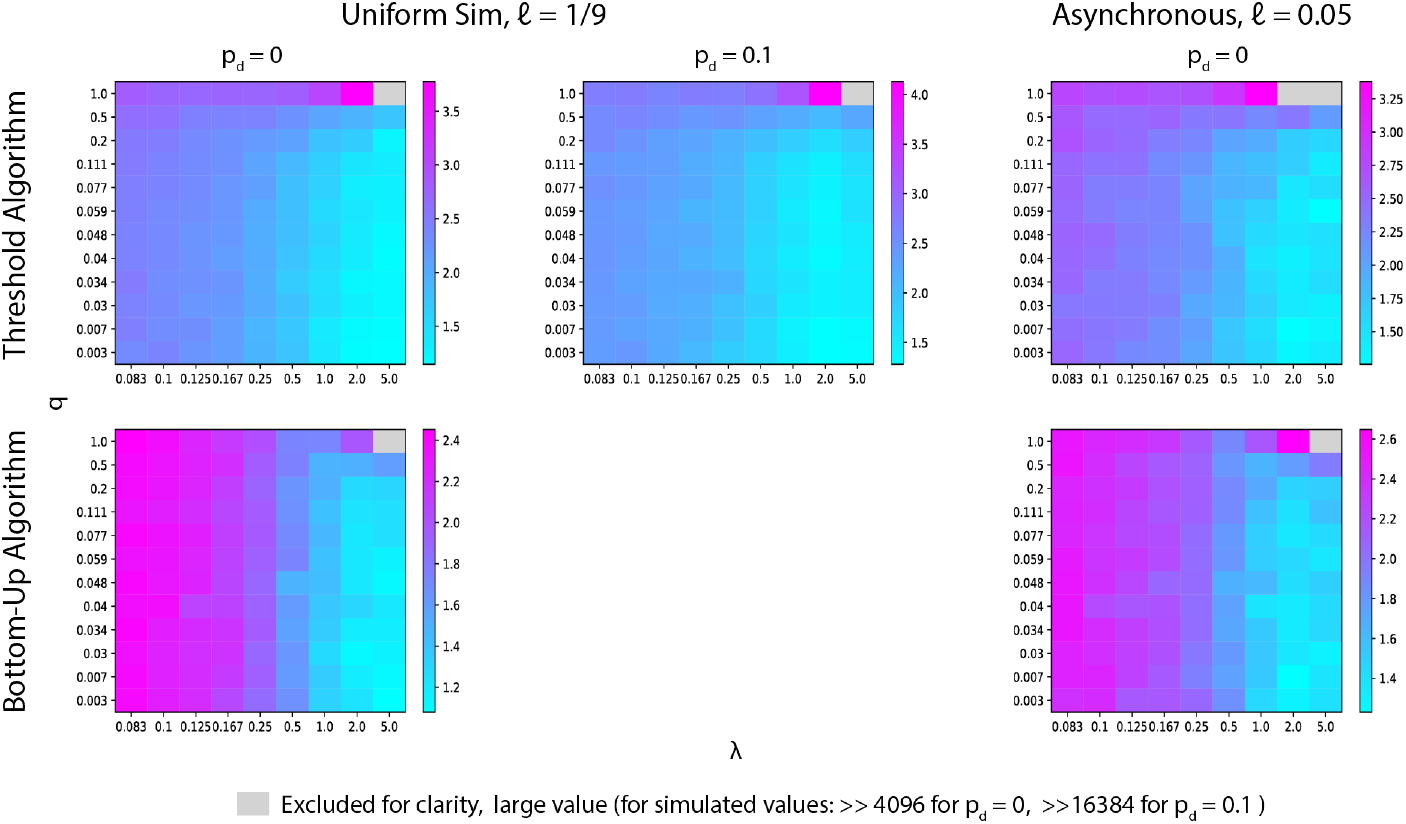
Triplets correct scores for both algorithms. Simulated trees with 256 leaves, *n* = 256. Entries are *log*_10_ scaled. Minimum *k* required for 0.9 probability of ≤ 0.95 proportion of 500 uniformly sampled triplets correctly reconstructed in simulation. Top row: Results for the Threshold Algorithm in the case of (From left to right) uniform edge length topology without missing data (*ℓ* = 1/9, *p_s_* = 0.1), uniform edge length topology with missing data (*ℓ* = 1/9, *p_s_* = 0.1), asynchronous topology (*ℓ* = 0.05). Bottom row: Results for the Bottom-Up Algorithm in the case of (From left to right) uniform edge length topology (*ℓ* = 1/9) and asynchronous topology (*ℓ* = 0.05).

Regarding the Threshold Algorithm, the reduction in the necessary *k* is potentially due to the fact that if triplets are sampled uniformly, most of them will have an LCA close to the root. We saw from the partial reconstruction results that the necessary *k* decreases across the board as *d*\* (the depth up to which triplets must be correct) decreases. We see also that large values of *λ* coincide with lower relative *k* in the triplets case than in the case of full reconstruction, just as in the case of *d*\* *<<* 1. Thus, we can treat the case of uniformly sampling triplets as similar to setting a low *d*\*. We see that both of these effects, the lower overall necessary *k* and the lower *k* for high values of *λ*, also hold true of the Bottom-Up Algorithm in the case of uniformly sampling triplets. This indicates that perhaps reconstruction of triplets that diverge at the top of the tree is easier and less affected by mutation saturation in the case of the Bottom-Up Algorithm as well.

